# Shifts in vernalization and phenology at the rear edge hold insight into adaptation of temperate plants to future milder winters

**DOI:** 10.1101/2024.08.16.608089

**Authors:** Antoine Perrier, Megan C. Turner, Laura F. Galloway

## Abstract

- Temperate plants often regulate reproduction through winter cues, such as vernalization, that may decrease under climate change. Studies of rear-edge populations, glacial relicts that persist in environments that have warmed since last glaciation, can provide insight into adaptive potential to milder winters.
- We studied how rear-edge populations have adapted to shorter winters and compared them to the rest of the range in the herb *Campanula americana*. Using citizen science, climate data and experimental climate manipulation, we characterize variation in vernalization requirements and reproductive phenology across the range and their potential climatic drivers.
- Rear-edge populations experienced little to no vernalization in nature. In climate manipulation experiments, these populations also had a reduced vernalization requirement, weaker response to changes in vernalization length, and flowered later compared to the rest of the range.
- Our results suggest shifts in phenology and its underlying regulation at the rear edge to compensate for unreliable vernalization cues. Thus, future milder winters may be less detrimental to these populations than more northern ones. Furthermore, our results showcase strong adaptive shifts at the rear edge of temperate plants’ ranges, highlighting the importance of these areas in studies of predicted future climates.

## Introduction

Organisms exposed to cyclical environmental changes often evolve mechanisms to sense these fluctuations and to time developmental shifts to occur under favorable conditions (Preston & Sandve, 2013). In temperate plants, vernalization, the prolonged exposure to non-lethal seasonal cold (Chouard, 1960), serves as an important cue so that key life-cycle transitions will occur after winter, e.g. vegetative growth and reproduction (Amasino, 2005). However, relying on such cues may be detrimental under rapidly changing environments. This is of particular concern in the context of ongoing global warming as it is expected to affect the strength and timing of seasonal temperatures experienced by natural and agricultural species (Willis *et al*., 2008; Blackman, 2017). Species requiring vernalization will likely be negatively affected by climate warming as shorter and milder winters become more common (Luedeling *et al*., 2011; Anderson, 2023). Such conditions are expected to lead to a reduction in reproduction (Padhye & Cameron, 2009; Liu *et al*., 2012; Satake *et al*., 2013) or shifts in reproductive phenology outside of optimal time-windows (Fitter & Fitter, 2002; Parmesan & Yohe, 2003; Love & Mazer, 2021; Faidiga *et al*., 2023; Geissler *et al*., 2023). It is therefore critical to understand response to seasonal cuing, and variation in this response, to determine possible consequences of expected warmer climates for temperate plant species.

Vernalization requirements and cue responses often vary within and between species in response to differences in winter conditions (Andrés & Coupland, 2012; Blackman, 2017; Preston & Fjellheim, 2022). This suggests that while these mechanisms are crucial to complete the life cycle, they vary enough to allow persistence across heterogeneous environments. Studies have explored variation in cueing reproduction across environments by testing for clines in vernalization requirements and phenological response to temperature gradients (Blackman, 2017). This body of work provides evidence of a reduction in vernalization requirements with increases in temperature (Wesselingh *et al*., 1994; Dijk *et al*., 1997; Boudry *et al*., 2002; Stinchcombe *et al*., 2005; Jokela *et al*., 2015). Reproductive phenology also varies over temperature gradients though a general pattern is less clear. For example, flowering is later with the later arrival of spring-like conditions at high latitudes than lower ones in *Arabidopsis thaliana* (Stinchcombe *et al*., 2004; Lempe *et al*., 2005). In contrast, earlier flowering occurs at higher latitudes in other species (e.g. Paccard et al. 2014; Vest and Sobel 2021). These studies reveal the potential for both the mechanism of environmental cuing as well as the pattern of reproductive phenology to evolve to track warming climates.

Populations at the warmer limits of temperate species ranges are particularly important for understanding adaptation to shorter winters. This part of the range, colloquially known as the “rear edge” (Hampe & Petit, 2005), often consists of relict populations that have persisted more or less in place at least since the last glacial maximum (LGM, c. 23 000–19 000 years ago; Hughes et al. 2013). Long-term persistence at the warmer range limit over multiple glacial cycles and under continuously warming climates after the LGM has likely resulted in adaptation to warmer climates (Hampe & Petit, 2005). With this history, the rear edge may provide insight into how species requiring vernalization adapt to mild winters, and which adaptations may be best suited to future warming climates. However, these populations are typically not sampled in studies investigating clines and therefore our knowledge of phenology and vernalization requirements in populations from the warmest habitats is limited (but see Vest and Sobel 2021).

Here we investigate whether rear-edge populations show patterns of differentiation consistent with adaptation to milder winters compared to populations from elsewhere in the range in the North American herb *Campanula americana.* This species requires vernalization, but occupies a wide latitudinal and temperature gradient (Fig. 1a), with ancestral rear-edge populations occurring in distinctively warmer climates in the southern parts of the range (Barnard-Kubow *et al*., 2015). To test for differentiation between the rear edge and the rest of the range, we first characterized differences in flowering phenology across latitude using range-wide observations in natural populations gathered from citizen science data, and then tested whether this variation reflects differences in winter or growing season climate (Observational Study). We also assessed genetic differences in phenology by testing for variation in flowering among populations from across latitudes raised under common greenhouse conditions (Experiment 1). We then evaluated plasticity in reproductive phenology in response to a range of natural growing season cues by raising populations sampled across latitudes in common gardens along a latitudinal gradient (Experiment 2). Finally, we tested differences in vernalization requirement and phenological plasticity in response to these winter cues by raising populations under experimentally manipulated vernalization length (Experiment 3). In total, these four approaches allow us to comprehensively test for differences in phenology and its regulation across the range of this species. Results of these experiments allow us to understand not only how rear-edge populations have adapted to their distinct habitats, but also shed light on how populations across the range vary in their response to milder winter, and ultimately the adaptation that may evolve to allow persistence under future conditions.

**Fig. 1:**
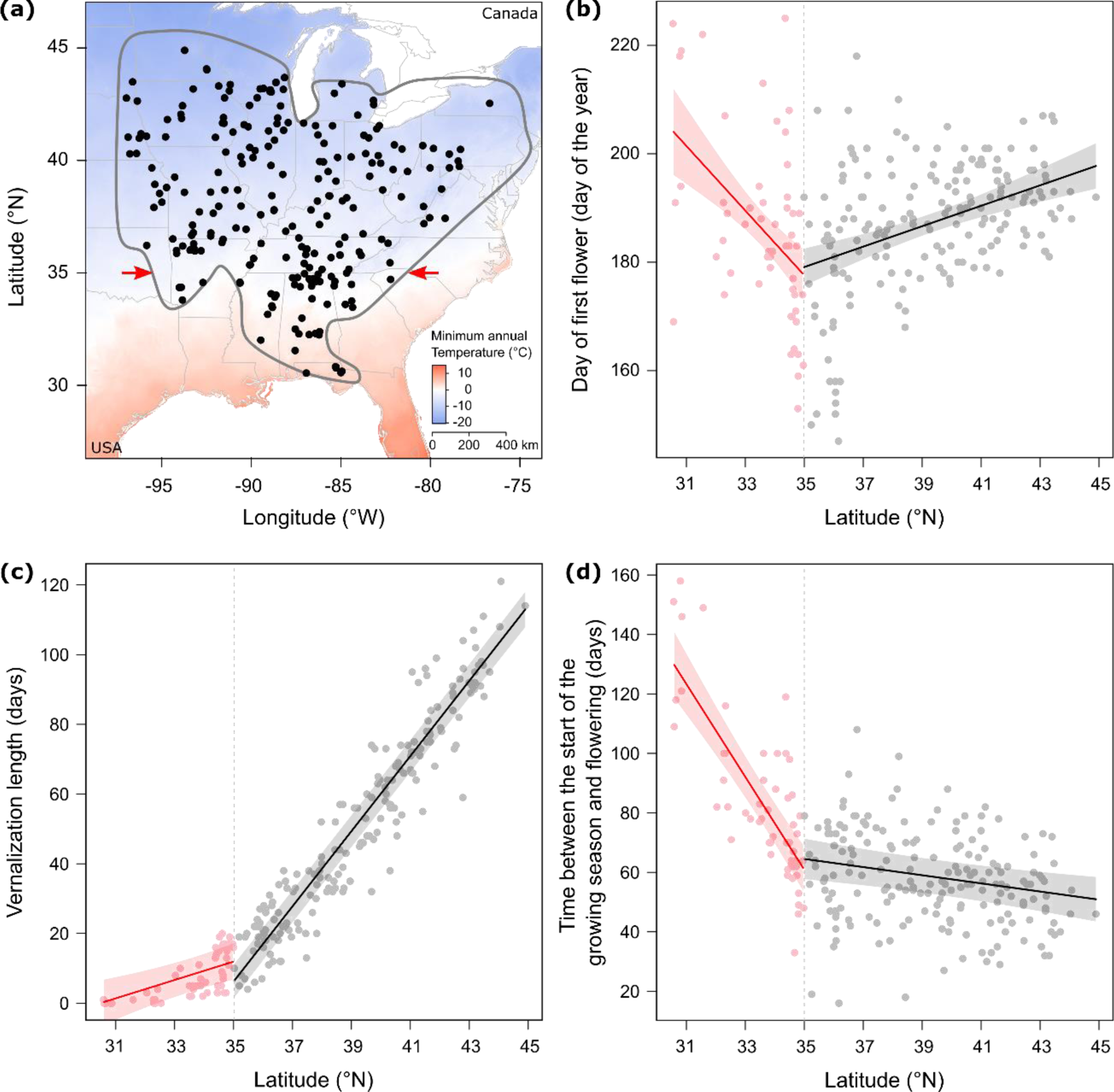
Variation in flowering phenology observed in nature and associated climatic factors across latitudes (Observational Study). (**a**) Location of *Campanula americana* observations (black dots) selected from iNaturalist 2018-2022. The range of the species is outlined with a gray line; red arrows represent the latitudinal delimitation of the rear edge. Shading indicates the latitudinal temperature gradient based on 30 year minimum temperature in January (1991-2020) from PRSIM climate data (https://prism.oregonstate.edu). (**b**) Day of first flower inferred for observations in (**a**). (**c**) The length of vernalization and (**d**) the time between the start of the growing season and flowering inferred for each observation from climate data. Colors represent whether observations occurred below (red) or above (black) the identified breakpoint (34.98° N, dashed vertical lines) in the latitudinal cline in day of first flower. Solid lines represent the significant model predicted slope of the relationship between each variable and latitude, respective to the breakpoint, with the 95% confidence interval indicated as shading. Test statistics reported in Table 1.

**Table 1:** Test of variation in the day of first flower, length of vernalization and time between the start of the growing season and flowering across the latitudinal range of *Campanula americana* relative to a predicted breakpoint.

| Dependent variable | Latitude (L) | | | | Breakpoint (BP) | | | L * BP | | | $R^2m$ | $R^2c$ |
| --- | --- | --- | --- | --- | --- | --- | --- | --- | --- | --- | --- | --- |
| | $\beta < BP$ | $\beta > BP$ | $\chi^2$ | | $\beta$ | $\chi^2$ | | $\beta$ | $\chi^2$ | | | |
| Day of first flower | -5.96 | 1.88 | 23.59 | *** | -273.10 | 39.52 | *** | 7.85 | 37.84 | *** | 0.18 | 0.18 |
| Length of vernalization | 2.63 | 10.75 | 13.57 | *** | -289.84 | 130.99 | *** | 8.12 | 119.59 | *** | 0.93 | 0.95 |
| Time between flowering and the start of the growing season | -15.58 | -1.37 | 107.87 | *** | -493.68 | 85.97 | *** | 14.25 | 82.81 | *** | 0.46 | 0.54 |

All dependent variables were assumed to follow Gaussian distributions. Each model was optimized with the *bobyqa* optimizer to improve convergence. Test statistics include the slope of each effect ( $\beta$ ), the $\chi^2$ -value, and the marginal and conditional $R^2$ of the model. For the effect of latitude, $\beta$ was estimated for the parts of the range below or above the predicted breakpoint (BP) in latitude (34.98 °N). \*\*\* $P < 0.001$ . Bonferroni corrections were applied for $P$ -values of each fixed effect for Day of first flower and Time between flowering and the start of the growing season as they are non-independent, but significance thresholds did not change. Results for random effects are not shown.

## Material and methods

### Study system

*Campanula americana* L. is a monocarpic herb found in open forests, clearings and forests margins across the eastern USA (Fig. 1a). Seeds typically germinate in spring or fall, vernalize over winter as rosettes and then bolt, elongating into a reproductive stalk. Following several months of growth, flower buds develop and the plants transition to flowering in mid-summer, with flowers opening progressively over several weeks. Most populations require at least six weeks of cold temperatures, estimated as 4° C, for successful reproduction (Baskin & Baskin, 1984; Etterson & Galloway, 2002), but the southernmost populations may not always require cold exposure to bolt (Kalisz and Wardle 1994). With this natural history, we define vernalization as any time period where rosettes are exposed to temperatures below 4° C.

*Campanula americana* occupies a large range, spanning ∼15 ° latitude over ∼1800km with a difference of ∼22 °C in minimum annual temperature (Fig. 1a; inferred from the PRISM database, https://prism.oregonstate.edu). The species comprises three geographically distinct genetic clades (Barnard-Kubow *et al*., 2015), but most populations (∼ 80%) belong to a clade found at low elevation (<600m) to the west of the Appalachian Mountains. Here we focus on this “Western” clade, which is characterized by a history of persistence in glacial refugia in the southern part of the current range during LGM, followed by postglacial range expansion (Barnard-Kubow *et al*., 2015; Koski *et al*., 2019; Prior *et al*., 2020). We define the “rear edge” as the lower latitudinal third of the range (below 35° N, Fig. 1a), where populations occur in former putative glacial refugia (Barnard-Kubow *et al*., 2015). This part of the range also has a climate defined as subtropical, characterized by mild winters and rare freezing events (American meteorological society, https://glossary.ametsoc.org/wiki/Subtropics), distinct from the continental temperate climate in more northern parts of the range.

### Observational Study: Variation in flowering phenology in natural populations

We first tested whether rear-edge populations in *C. americana* differ from the rest of the range based on flowering phenology observed in natural populations across latitudes, and then explore associations between phenology and both winter and growing season climates.

#### Inference of flowering phenology in natural populations

We inferred flowering phenology of natural populations from the publicly available repository iNaturalist (https://www.inaturalist.org. Accessed 23/02/2023). We extracted all 10,250 records of *C. americana* observed between 2018 and 2022 (earlier records did not cover the distribution well). This set was reduced to 335 observations following pruning to select for low elevation populations, reduce duplicates and oversampling close to cities, and reduce bias of fewer occurrences at the rear edge (Methods S1, Fig. S1). We inferred the day of first flower (recorded as day of the year) based on the photograph provided with each observation (Table S1, Fig. S2), resulting in a final dataset of 238 observations (Fig. 1a).

#### Climatic variables

For each observation, we obtained daily mean, maximum, and minimum temperature and daily precipitation for the year of observation and the year before from the PRISM database (accessed 01/09/2023) using the *prism* package (Edmund & Bell, 2015) in R (R Core team, 2024). We then generated 17 variables from this data that capture the climate at key life cycle stages to test associations of phenology with climate (Table S2a).

#### Statistical analysis

We initially explored how the day of first flower varied across latitude. To do this, we fit three mixed-effect models with latitude as a continuous fixed effect and the year of observation as a random effect, and tested which model best described variation in phenology through model selection. The first model tested for a linear relationship between the day of first flower and latitude, the second tested a second-degree quadratic relationship, and the third model tested a piecewise linear relationship with one breakpoint. The latter two allow clines with latitude to vary across the range. All analyses were performed in *R*. Model parametrizations are provided in Methods S2a. Model fits were compared using the corrected Akaike’s information criterion (AICc, Sugiura 1978) using *MuMin* (Bartoń, 2023). From this we identified a breakpoint in the relationship between the day of first flower and latitude at ∼35 °N (see Results) that we incorporated into the design of subsequent experiments.

We then assessed how parts of the range that differ in patterns of day of first flower also differ in climate. We first tested which of the 17 climatic variables generated above best discriminated between the part of the range below or above the breakpoint in latitude by performing a discriminant analysis of principal components (DAPC) using the *dapc* function in *adegent* (Jombart, 2008; Jombart *et al*., 2010). Climatic variables, especially those related to temperature, are often correlated with latitude. We performed a DAPC because it reduces correlation between variables before the discriminant analysis by transforming them using principal component analyses (PCA). DAPC was performed on all climate variables (8 PC retained) with the grouping factor of whether observations were located at latitudes above (group 1) or below (group 2) the predicted breakpoint, effectively identifying variables that had non-linear relationships with latitude. We then evaluated how the climatic variables that contributed the most to the discriminant function (>0.05) varied across the range to understand their potential influence on patterns of flowering. To do this, we tested the variables in mixed effect models that included the latitude of each observation as a continuous fixed effect, range position of each observation relative to the estimated breakpoint as a categorical effect, and their interaction. Year of observation was included as a random effect (model in Methods S2b). We also tested the effect of latitude and range position on the day of first flower using the same model structure for comparison. In these models, variables were analyzed assuming a normal distribution. For each model, we tested and confirmed model assumptions.

### Experiment 1: Variation in reproductive phenology in a common environment

We tested for genetic differences in flowering phenology between *C. americana*’s rear edge and the rest of the range by raising populations from across latitudes under a common environment in a greenhouse experiment.

#### Experimental design

We raised two cohorts of *C. americana* populations that cover a similar latitudinal breadth below (∼4.4° N) and above (∼6.6° N) the breakpoint identified in the Observational Study. For the first cohort, we collected seeds in 23 natural populations (Fig. 2a, Table S3) in late summer of 2020 and 2021 and stored them by maternal plant at 4 °C under dark and dry conditions. For the second cohort, we used greenhouse-reared seeds from 25 populations (Methods S3), 20 from the first cohort and five additional populations to yield better sampling across latitudes (Fig. 2a). Greenhouse produced seeds have less variation due to maternal environment. In this species, maternal effects mostly affect germination and are thus not of concern for reproductive phenology (Galloway, 2005; Galloway & Burgess 2009). We raised 544 seedlings in the first cohort (Fall 2021), and 535 in the second cohort (Fall 2022) in similar conditions (Methods S3). Conditions were designed to mimic the natural growth cycle of the species, starting with germination and vegetative growth in growth chambers simulating fall, followed by vernalization in a cold room simulating winter, and then bolting and reproduction in a greenhouse with conditions simulating summer.

**Fig. 2:**
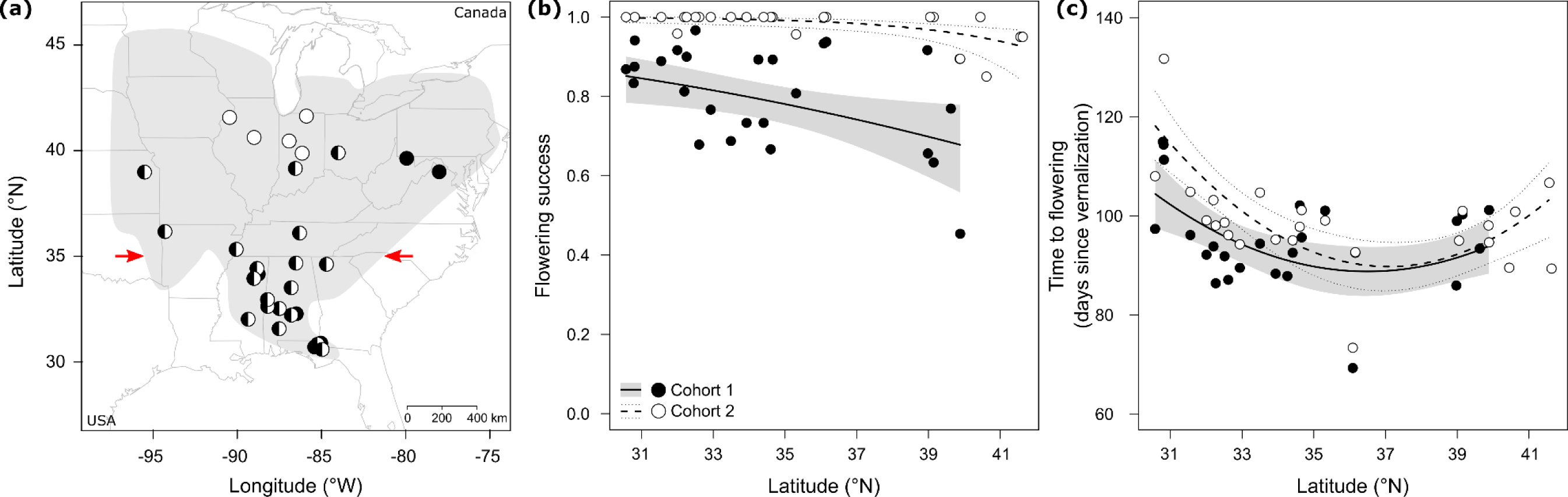
Variation in reproductive phenology in a common environment (Exp. 1). (**a**) Location of *Campanula americana* populations (dots). The range of the species is shaded in gray; red arrows represent the latitudinal delimitation of the rear edge. Mean flowering success (**b**) and time to flowering (**c**) for two cohorts of populations (dots) raised in a common controlled environment. Lines represent the significant model-predicted relationship between traits estimated at the individual level and population latitude, with the 95% confidence interval indicated as shading (cohort 1) or dotted lines (cohort 2). Test statistics reported in Table 2 and Table S8.

**Table 2:**
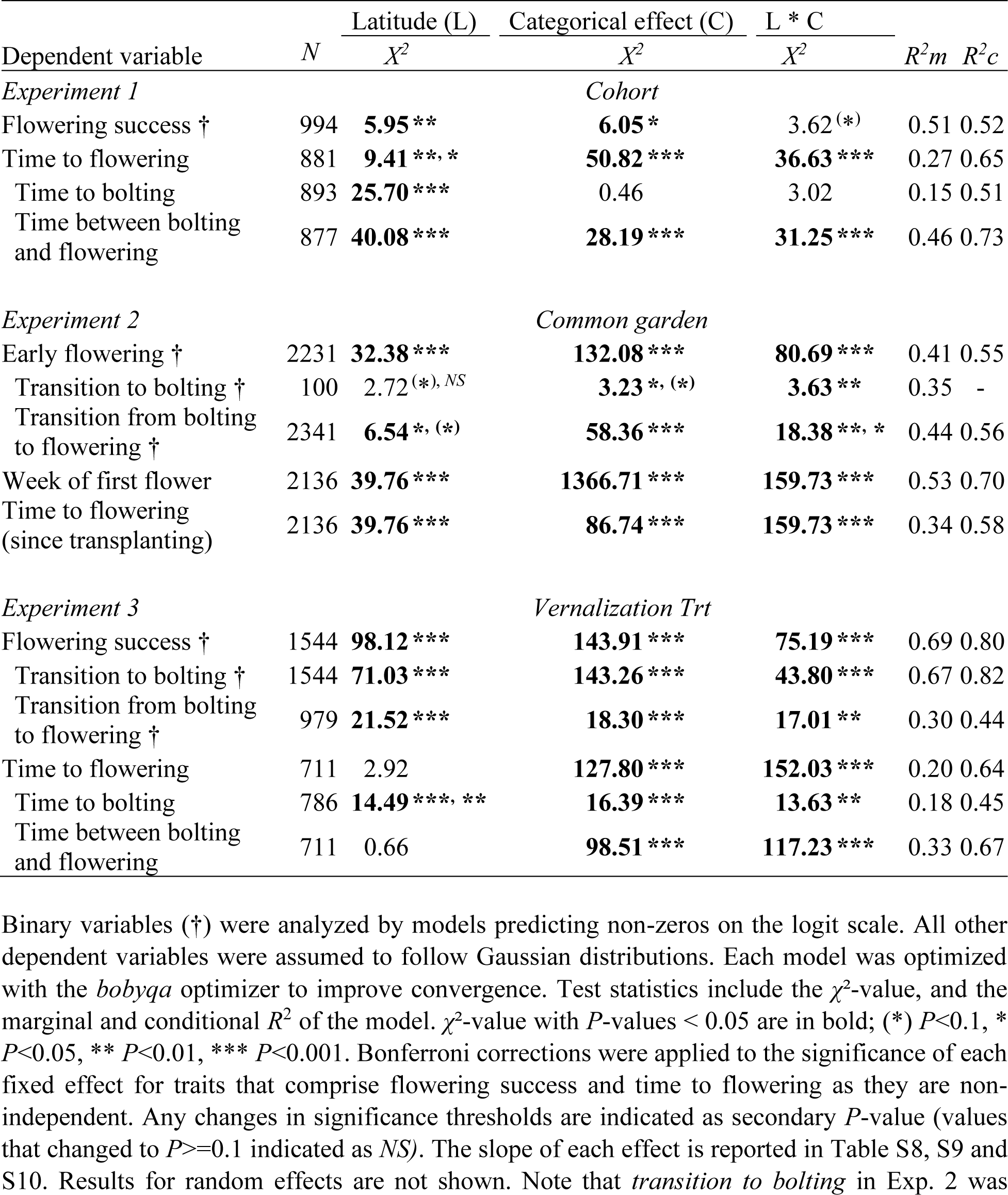
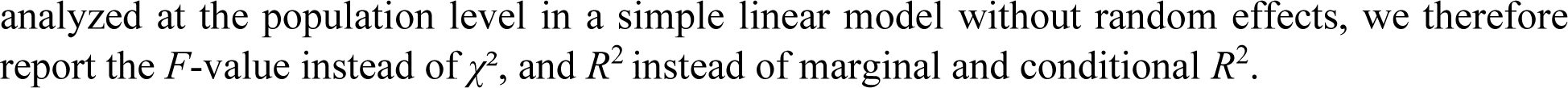
Test of variation in reproductive phenology of *Campanula americana* across latitudes in a common greenhouse environment (Exp. 1), common gardens across a latitudinal gradient (Exp. 2) and in response to different length of vernalization treatments (Exp. 3).

#### Recording of traits and statistical analysis

We recorded the date of bolting (presence of a stem) twice a week for the first cohort and weekly for the second. We recorded the date of first flower three times a week for both cohorts. Most plants that bolted also flowered (>98%).

Based on this data, we generated two main traits related to reproduction initiation and phenology (Table S4a): *flowering success* (binary, production of at least 1 flower) and *time to flowering* (days from vernalization to flowering). We additionally decomposed *time to flowering* into two components *time to bolting* and *time between bolting and flowering* to better understand which developmental stage underlies the overall pattern in phenology. Each trait was analyzed at the individual level in mixed effect models in *R*, with population *latitude* as a continuous fixed effect, *cohort* a categorical effect, and their interaction. Population and seed family nested within population were random effects (models in Methods S4). Here and for each subsequent experiment, binary traits were analyzed assuming a binomial distribution, and timing variables were analyzed assuming a normal distribution. For each model, we tested and confirmed model assumptions. Similar to the Observational Study, we initially tested whether latitude was best described as a linear relationship or a second-degree quadratic relationship for each trait. We chose to test a quadratic relationship instead of a breakpoint for ease of interpretation, as the breakpoint approach would require integrating additional two-way and three-way interactions. Comparison of model performance was based on AICc. We report results only for the best performing model (all results Table S5). For this model, we performed a post-hoc test to estimate differences in traits between cohorts, and differences in the effect of latitude on traits between cohorts, using the package *emmeans* (Lenth, 2019).

### Experiment 2: Variation in reproductive phenology in a transplant experiment

We tested for differences in phenology in response to spring and summer conditions between the rear edge and the rest of the range by raising populations from across the range in common gardens established on a latitudinal gradient.

#### Experimental design

We selected 25 populations with similar distribution as Exp. 1 (Fig 3a, Table S3), and approx. 15 seed families per population (Table S3), to be raised in five common gardens. We used greenhouse-reared seeds for 20 of the populations and field-collected seeds for five. For each garden, we sowed two seeds in each of two pots per seed family (three seeds for field-collected seeds, one if few seeds), thinning to one plant per pot after germination. This gave 2 individuals/family x 15 families/population x 25 populations = 750 individuals per garden, 3750 total. Seeds were planted in fall 2022, germinated for six weeks in growth chambers and vernalized for seven weeks in a cold room (Methods S3).

**Fig. 3:**
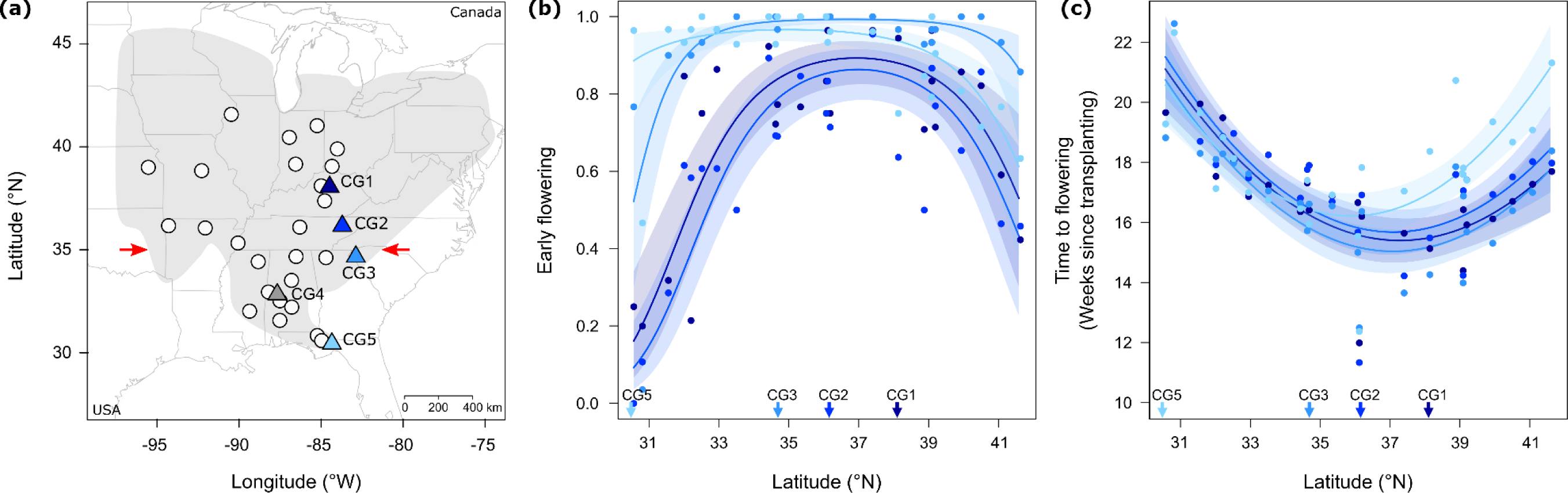
Range-wide variation in reproductive phenology in common gardens (Exp. 2). (**a**) Location of *Campanula americana* populations and common gardens. The range of the species is shaded in gray; red arrows representing the latitudinal delimitation of the rear edge. Populations are indicated by white dots, and common gardens by colored triangles. CG4 (gray) was not included in the analyses because of low survival. Proportion of early flowering (**b**) and mean time to flowering since transplant (**c**) were recorded for populations (dots) sampled across the range and raised in common gardens across a latitudinal gradient. Colors distinguish the different gardens as depicted in (A), and their latitude is indicated with arrows along the x-axis. Lines represent the significant model-predicted relationship for each garden between traits estimated at the individual level and home latitude of the population, with the 95% confidence interval indicated by shading. Test statistics reported in Table 2 and Tables S9.

Seedlings were acclimated to outdoor conditions for one week (Charlottesville, VA, USA), and then transplanted to one of five common garden sites (CG, Fig. 3a, Table S6): Lexington KY (CG1, 38.1° N), Blaine TN (CG2, 36.2° N), Clemson SC (CG3, 34.7° N), Akron AL (CG4, 32.9° N), and Tallahassee FL (CG5, 30.5° N). These sites span an 850 km latitudinal gradient in roughly 2° steps and reflect the temperature gradient within the sampled range. Transplant occurred in spring 2023, starting with CG5 and ending with CG1, to reflect the later start of the growing season at higher latitudes. The transplant date for each site was the date when mean daily temperature was consistently above 10°C in spring, estimated using PRISM daily mean temperature 2018 to 2022. Within each site, all seedlings were transplanted in one or two days. Sowing was timed for each site such that vernalization ended one week before the projected transplant date, ensuring seedlings were transplanted at the same life stage across the experiment.

Garden locations were chosen to mimic the natural habitat of the species. In each site, individuals were transplanted in 30 spatial blocks of 25 individuals, with one individual per population per block. Sites were raked to reduce initial competition. Individuals were then transplanted into the exposed ground with the substrate they were raised in and watered once. Each site was surrounded by a 2m high fence to exclude large herbivores. No further intervention was performed. One data logger (iButton®, Maxim Integrated Products, Inc) was placed in each site in the shade, 1.5m above ground, to monitor air temperature every hour for the length of the experiment. This data revealed that plants experienced additional days of vernalization after transplant (Table S6a), but less so in the southernmost garden. The Experiment was stopped 153 to 170 days after transplant (Table S6b), when most plants had reached fruit maturation, but seeds were not yet dispersing. Unfortunately, plants in CG4 experienced very low survival (<10%), so this site was removed from analysis.

#### Recording of trait and statistical analysis

Data was collected three times, about a month apart, in the three northern sites (GC1-GC3), and only two times for the southern site (GC5). During each data collection visit, we recorded survival, transition to bolting and to flowering (here production of at least one bud) and inferred the time of first flowering (Methods S5).

Based on this data, we generated the plant-level binary trait *early flowering,* defined as plants with at least one open flower at the first data collection visit, to provide a “snapshot” of phenology (Table S4b). At the end of the experiment, we assessed two binary components of flowering success, whether plants *transition to bolting* and *transition from bolting to flowering*. These were evaluated to determine the sensitivity of different components of reproduction to local environmental cues. We assessed flowering phenology by estimating the *Week of first flower* (since January 1^st^), as well as *time to flowering since transplant* (Weeks from transplant) to take in account the differences in transplant date between gardens. The effects of latitude (of population origin), common garden (categorical, four sites) and their interaction on each of these traits was analyzed in mixed effect models (Methods S4). Preliminarily analysis tested whether latitude was best described by a linear or a quadratic relationship, and results are only reported for the best model (Table S5). If models performed equally well, the quadratic one was retained for ease of comparison among traits. For the best models, we performed post-hoc tests to assess differences in traits between each garden, and differences in the effect of latitude on traits between gardens, using the package *emmeans* (Lenth, 2019).

### Experiment 3: Variation in vernalization requirements and response across the range

We tested for differentiation in vernalization requirement and in phenological response to vernalization length between the rear edge and the rest of the range by subjecting populations from across latitudes to experimentally manipulated vernalization length.

#### Experimental design

We selected 12 populations with greenhouse-reared seeds, representing the same latitudinal range as Exp. 1 (Fig. 4a, Table S3). We selected approx. 15 families per population (Table S3), and sowed two seeds in each of nine pots per seed family, resulting in 135 individuals per population after thinning to one seedling per pot. Pots were then randomly assigned to one of three vernalization treatments (see below), resulting in 3 individuals/family x 15 families/population x 12 populations = 540 individuals per treatment (45 per population/treatment), 1620 total. Seedlings were raised from germination in fall 2022 to flowering in 2023 (Methods S3), but with the difference that treatments differed in vernalization length, lasting either six, four or two weeks. Sowing was timed such that all three treatments finished vernalization at the same time (Fig. S3). The six weeks treatment represents vernalization conditions experienced by a mid-latitude population (Baskin & Baskin, 1984), that induces vernalization across the range and is slightly shorter than that used in prior controlled environment studies (e.g. Prendeville et al. 2013). The four and two week treatments represent even time intervals between zero and six weeks. A “no vernalization” treatment was not tested because as at least some vernalization is required for flowering to occur in most populations (Kalisz & Wardle, 1994).

**Fig. 4:**
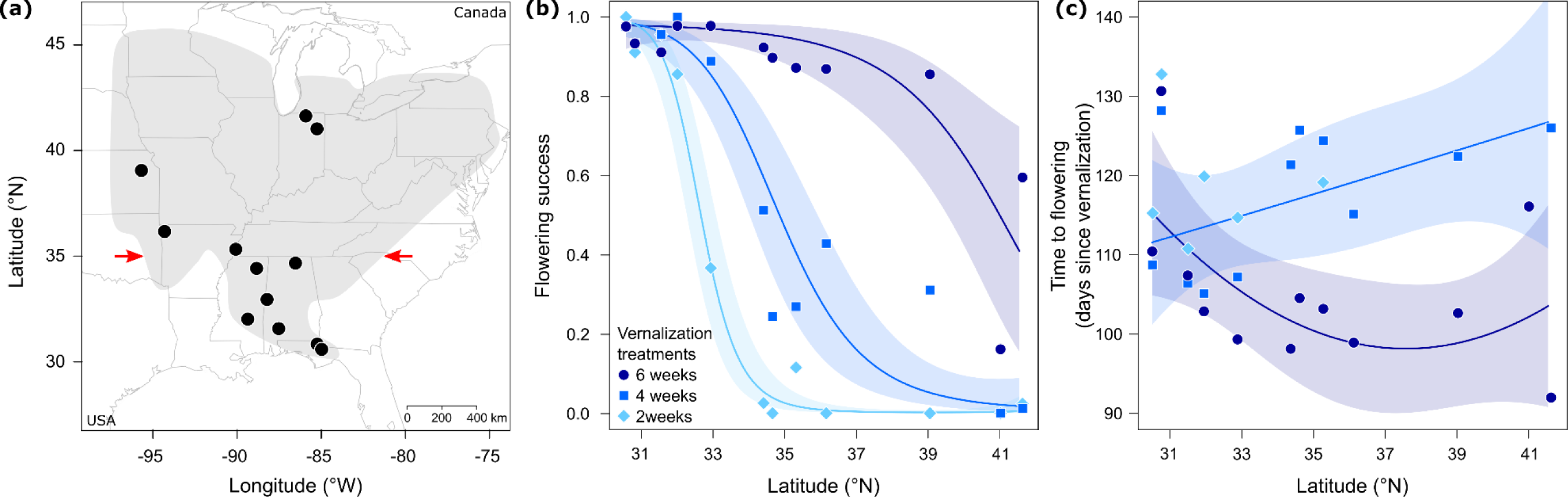
Effect of vernalization length on range-wide variation in flowering success and phenology (Exp. 3). (**a**) Location of *Campanula americana* populations (dots). The range of the species is shaded in gray; red arrows represent the latitudinal delimitation of the rear edge. Flowering success (**b**), and mean time to flowering (**c**) for populations (dots) sampled across the range and exposed to three vernalization treatments (six, four or two weeks) in controlled conditions. For time to flowering, the 2-week treatment was excluded from the analysis as too few individuals flowered (population means based on at least five individuals shown for comparison). Lines represent the significant model-predicted relationship between each trait estimated at the individual level and population latitude in each treatment, with the 95% confidence interval indicated as shading. Test statistics reported in Table 2 and Tables S10.

#### Recording of traits and statistical analysis

We recorded survival, bolting and flowering three times a week once plants were moved to the greenhouse. Bolting was recorded for 52 days after vernalization and flowering for 141. We then estimated *flowering success,* and its two components *transition to bolting* and *transition from bolting to flowering*, as well as *time to flowering* and its two components *time to bolting*, and *time between bolting and flowering* (Table S4a). Similar to Exp. 1, the effects of latitude, vernalization treatment (categorical with three levels) and their interaction on each of these traits was analyzed in mixed effect models (Methods S4). For the three timing traits, the two-week treatment was excluded as few individuals bolted. Preliminarily analysis tested whether latitude was best described by a linear or a quadratic relationship (Table S5) and results are reported for the best model. For the best models, we performed post-hoc tests to assess differences in traits between each vernalization treatment, and differences in the effect of latitude on traits between treatments, using the package *emmeans* (Lenth, 2019).

## Results

### Observational Study: Variation in flowering phenology in natural populations

In natural populations, the relationship between the day of first flower and latitude was best described as a quadratic or piecewise relationship (Table S7, Fig. S4), with both models performing equally well (ΔAICc < 2). The earliest flowering was at 36.64° N and 34.98° N respectively, with flowering at increasingly later dates towards lower and higher latitudes. Subsequent analyzes were parameterized with the breakpoint from the piecewise model because it matched best our *a priori* delimitation of the rear edge (∼35 °N), though use of the minimum from the quadratic model yielded similar results (not shown). Climates of populations north and south of this breakpoint were best discriminated by the length of vernalization (LVerna) as well as the time between the start of the growing season and flowering (FlowGS, Table S2b). The other climate variables made negligeable contributions to discriminating between the regions of the species range above and below the breakpoint.

The day of first flower and the two climate variables (length of vernalization, time between the start of the growing season and flowering) showed strong clines across latitudes, with the slope depending on their position relative to the breakpoint (L*BP, Table 1). Flowering was earliest at the breakpoint, with much later flowering per unit of latitude toward low latitudes (26 days over ∼4.5 °) than towards high latitudes (18 days over ∼9.9 °, Table 1, Fig. 1b). The length of vernalization increased with latitude (Table 1, Fig. 1c), but days of vernalization accumulated more rapidly north of the breakpoint than to the south, where vernalization was limited. The time between the start of the growing season and flowering was substantially shorter and relatively insensitive to latitude to the north (65 to 51 days predicted), whereas it increased sharply to the south of the breakpoint (62 to 130 days predicted; Table 1, Fig. 1d).

### Experiment 1: Variation in reproductive phenology in a common environment

Rear-edge populations showed differentiation in reproductive phenology from the rest of the range when raised in a common controlled environment. Populations from low latitudes had greater flowering success but flowered later relative to those in the rest of the range (Table 2, Fig. 2, Table S8). Populations from the highest latitudes also flowered later than central populations, but this pattern was weaker than the low latitude populations. Later flowering was due to a delay in time to bolting in northern populations, but a longer developmental period between bolting and the opening of the first flower in southern populations (Table 2, Table S8, Fig. S5). Patterns across latitudes were consistent between cohorts, though a greater proportion of plants flowered in the second cohort, for which only the northernmost populations showed reduced success.

### Experiment 2: Variation in reproductive phenology in a transplant experiment

Rear-edge populations showed strong differentiation from the rest of the range in the common gardens due to delayed reproduction relative to those from more central sites. Southern populations had fewer early flowering plants (except in CG5, Table 2, Fig. 3b, Tables S9). These populations also flowered later, both in week of first flower and time to flowering since transplanting (Table 2, Fig. 3c, Tables S9, Fig S6c). Southern populations were less likely to bolt in the two northern gardens, but most individuals that bolted also flowered (Table 2, Tables S9, Fig S6a, b).

Populations from high latitudes were also less likely to flower early and had slightly later flowering time relative to central populations in all gardens, but delays in phenology were weaker than low latitude populations (except in CG5). Plants from northern populations only bolted in large numbers in the central garden, though most plants that bolted also flowered (Table 2, Tables S9, Fig S6a, b).

Patterns of phenology among southern populations were generally consistent across garden sites despite differences in spring and summer climate (Fig. 3). In contrast, northern populations exhibited plasticity in flowering phenology across gardens. For example, in the most southern garden (CG5), northern populations showed a stronger delay in phenology relative to central populations than in any of the other sites (Table 2, Fig. 3, Tables S9, Fig S6).

Variation in phenology across gardens was generally linked to northern gardens having been transplanted later than southern gardens. The early flowering proportion was significantly smaller in northern sites compared to southern sites (except for CG1 vs CG2; Fig. 3b, Table S9b), indicating a delay in flowering in northern sites. In line, the week of first flower generally occurred later in northern sites compared to southern sites (except for CG1 vs CG2; Table 2, Table S9b, Fig. S6c). However, these differences were almost eliminated when flowering was measured relative to transplant date (Fig. 3c, Table S9b).

### Experiment 3: Variation in vernalization requirements and response across the range

Vernalization requirements of rear-edge populations were strongly differentiated from the rest of the range, characterized by a reduced need for vernalization and smaller plastic response to variation in vernalization length. In keeping with this pattern, there was a decrease in flowering success in populations from higher latitudes that was exacerbated when vernalization was reduced (Table 2, Fig. 4b, Table S10). In northern populations, a reduction in vernalization from six weeks to two weeks yielded a large reduction in flowering success. In contrast, southernmost populations had high flowering success regardless of vernalization treatment. Most of this variation was due to variation in transition to bolting among populations (Table 2, Tables S10, Fig. S7a).

Differentiation in response to vernalization was also observed for phenology, with rear-edge populations again showing smaller plastic response to variation in vernalization. In the six-week treatment, populations from low latitudes flowered later compared to more central populations (Fig. 4c). Reducing vernalization led to a general delay in time to flowering except for in the southernmost populations which changed little (Fig. 4c, Tables S10). Differences in time to flowering across treatments were mainly due to changes in the time between bolting and flowering, rather than changes in time to bolting (Table 2, Tables S10, Fig. S7).

## Discussion

Understanding how relict rear-edge populations at the warmer range limits of temperate species differ from younger populations at higher latitudes can help us predict how populations across the range may respond and adapt to future climates. We found that in the North American herb *Campanula americana,* rear-edge populations occurring in the lower latitudinal part of the range have evolved to rely less on winter cues to regulate phenology compared to the rest of the range. This suggests that critical life cycle mechanisms, such as the regulation of phenology, may evolve rapidly to adapt to changing environments. In addition, contrary to classic expectations, we found the warmer range limit may be less affected by climate change’s mild winters compared to higher latitude regions of the range. Finally, our results suggest that northern populations will suffer losses in reproduction with a reduction in winter cues although, over time, they may adapt to these changes similarly to the rear edge.

### Natural rear-edge populations differ in phenology from the rest of the range

In natural *C. americana* populations, timing of flowering had opposing clines across latitudes (Observational Study, Fig. 1b). The breakpoint of the clines was ∼35 °N, coinciding with our *a priori* definition of the rear-edge (below) relative to the rest of the range (above). The day of first flower was the earliest at the breakpoint, becoming progressively later towards higher and lower latitudes. Previous work has found latitudinal clines in flowering are most often continuous in temperate plants (e.g. Stinchcombe et al. 2004). These clines are thought to reflect continuous differences in cues such as photoperiod or vernalization used to signal the onset of favorable growing seasons (Preston & Sandve, 2013; Blackman, 2017; Preston & Fjellheim, 2022). However, rear-edge populations have typically been neglected in such studies, so we don’t know how common the opposing clines seen here are. Opposing clines suggest a divergence in the response to seasonal cues, or a change in the cues used to regulate phenology between the rear edge and the rest of the range.

### Most of the range shows adaptation to winter conditions typical for temperate plants

We found that differences in the day of first flower between rear-edge populations and the rest of the range were associated with differences in the length of vernalization (Observational Study), suggesting vernalization may serve as the environmental cue for flowering. In nature, populations occurring above 35 °N experience increasing winter length towards higher latitudes (Fig. 1c). These longer winters were associated with an increase in vernalization required for reproduction in both greenhouse experiments (Exp. 1, 3). Fewer plants flowered from high-latitude populations exposed to short vernalization periods, and this response was strongest in the northernmost populations which had a complete loss in reproduction under limited vernalization. A longer vernalization requirement in high latitudes is common for temperate plant taxa (Wesselingh *et al*., 1994; Boudry *et al*., 2002; Jokela *et al*., 2015), and may reflect an adaptation to prevent premature flowering in areas with long winters and detrimental early spring frost events (Inouye, 2000).

Differentiation in the day of first flower across latitudes was also explained by the time between flowering and the start of the growing season. Above 35 °N, flowering in nature occurred somewhat earlier relative to the onset of the growing season with increasing latitudes (ca. 14 days earlier across ca. 10° of latitude to the north, Observational Study, Fig. 1d). This is also common among temperate plants (e.g. Paccard et al. 2014; Vest and Sobel 2021), and is thought to be associated with adaptation to complete reproduction under shorter growing seasons (Griffith & Watson, 2005). A similar shift in phenology was reported along an elevational gradient in *C. americana*, where high elevation populations with shorter growing seasons had accelerated reproduction compared to low elevation populations (Haggerty & Galloway, 2011).

In the manipulative experiments, populations from locations north of the rear edge exhibited plasticity in flowering phenology in response to vernalization length. While generally flowering slightly later than central populations, they flowered even later when vernalization was short (Exp. 3, Fig. 4c) or when raised in southern climates (Exp. 2, Fig. 3c), with northernmost populations exhibiting the strongest delay. This plasticity of later flowering under shorter vernalization may simply be a passive byproduct of inadequate cuing. Specifically, limited vernalization may partially cue the onset of reproduction, therefore resulting in a longer transition to reproduction. The greater plasticity in timing of flowering in northern populations may thus be due to longer vernalization requirements, increasing the mismatch between required and experimental vernalization length. Indeed, the shortfall for vernalization for northern populations in the manipulative experiments may have masked the earlier flowering relative to the start of the growing season observed in nature. Alternatively, greater plasticity in the northernmost populations could have evolved to modulate reproductive phenology in response to thaws in late winter, extending the period prior to initiating reproduction in short winters and thereby reducing negative effects of these “false springs”. In other studies, sensitivity to vernalization length may depend on local environment but is inconsistent, with populations from both warmer (e.g. Stinchcombe et al. 2005; Vest and Sobel 2021) and colder environments (Boudry *et al*., 2002; Jokela *et al*., 2015; Landoni *et al*., 2022) more sensitive to vernalization length. Greater sensitivity has been hypothesized to allow plants to modulate phenology relative to yearly variation in growing season length (Boudry *et al*., 2002; Vest & Sobel, 2021; Landoni *et al*., 2022).

### Winter cue requirements and responses are shifted at the rear-edge where winters are milder

Rear-edge populations showed distinct patterns of reproductive phenology compared to the rest of the range, suggesting differences in adaptation in response to milder, shorter winters. In nature, rear-edge populations experience less vernalization, with some not experiencing any vernalization. Southern *C. americana* populations reflect this climate difference by having reduced vernalization requirements, including southernmost populations which achieve a high percentage of flowering (>90%) even under very limited vernalization (Exp. 3, Fig. 4b; Kalisz & Wardle, 1994). Reduced vernalization requirements at the rear edge likely evolved in response to local shorter, warmer winters where vernalization was an unreliable cue. Such patterns have been found in other plants, in some cases leading up to a complete loss of a vernalization requirement (Wesselingh *et al*., 1994; Dijk *et al*., 1997; Boudry *et al*., 2002; Stinchcombe *et al*., 2005; Jokela *et al*., 2015).

Flowering in natural rear-edge populations was later relative to the onset of the growing season towards lower latitudes, with an almost ten-fold steeper cline in flowering time relative to the rest of the range. Flowering was also much later in rear-edge populations under common greenhouse conditions (Exp. 1, Fig. 2c), as well as in common gardens (Exp. 2, Fig. 3c). Such a dramatic difference in phenology between populations along temperature gradients has rarely been documented in temperate plants. The exception, wild beets, have later flowering in southern populations and variation in vernalization along a latitudinal gradient (Dijk *et al*., 1997), similar to *C. americana*. While it is possible that other environmental factors such as day length cue differences in phenology (Blackman, 2017), this seems unlikely for rear-edge populations in our study. Phenology in southernmost populations was not affected by variation in winter length (Exp. 3, Fig. 4c), growing season climate or planting date (Exp. 2, Fig. 3c), suggesting that the time to flowering is strongly genetically determined in these populations rather than relying on an alternative environmental cue to vernalization.

Together, our results suggest a dramatic shift in the regulation of flowering phenology at the rear edge, with less reliance on vernalization to initiate and time flowering. Instead, phenology in these populations is late and environmentally insensitive, contrasting with the plasticity in flowering time observed in northern populations. This evolved delay may represent an adaptation to compensate for the extremely early start of the growing season, thus allowing reproduction to track the warmest period of the year. A previous artificial selection study revealed trade-offs between early flowering and reproductive fitness in *C. americana* (Burgess *et al*., 2007), supporting an evolutionary advantage to delay flowering when the growing season length does not strongly limit reproductive output.

### The rear edge will differ in response to climate change compared to the rest of the range

Our study provides insight into how temperate plants may vary in their response to climate change across their range. Warming climates are often predicted to result in range shifts towards higher latitudes and elevations through colonization at the colder edge and extinctions at the warmer edge (Thomas *et al*., 2004; Hampe & Petit, 2005; Lenoir & Svenning, 2013). In temperate plants that rely on vernalization to cue reproduction, one may expect that populations at the warmer range limits will be especially sensitive to climate change as warming winters may not meet vernalization requirements, thus preventing the transition to reproduction (*e.g.* Padhye and Cameron 2009; Satake et al. 2013). However, in contrast to this prediction, we found that rear-edge populations of *C. americana* are likely to be the least affected by a reduction in winter length. Rather, populations at mid- and high latitudes may be at risk because their sensitivity in the initiation and timing of reproduction in response to winter length may lead to a fitness loss under milder winters. Reduced sensitivity at the rear edge may be common across temperate plants. Lower latitude populations often have a reduced vernalization requirement (Wesselingh *et al*., 1994; Dijk *et al*., 1997; Boudry *et al*., 2002; Stinchcombe *et al*., 2005; Jokela *et al*., 2015). They also show strong local adaptation (Bontrager *et al*., 2021), likely facilitated by a history of long term exposure to increasingly warm climates. This result contributes to a growing body of literature questioning the expected population decline under climate change at the rear edge (Vilà-Cabrera *et al*., 2019).

### Postglacial colonization history drives divergence in adaptation to changing climates

Our study highlights the importance of studying the effects of warming climates in a phylogeographic context, as the divergence between northern and southern parts of the range may also reflect different histories of exposure to climate change. The ∼35°N breakpoint in the day of first flower coincides with the transition between populations persisting in former glacial refugia in southern parts of the range, *i.e.* the rear edge, and more northern populations that have established after last glacial maximum (Barnard-Kubow et al. 2015; Koski et al. 2019; Prior et al. 2020). Post-glacial colonization in temperate plants often results from populations establishing in newly suitable habitats, tracking the gradual retreat of ice sheets as Earths’ global temperatures rose during the Holocene (Hewitt, 2000, 2004). In this context, *C. americana* populations at higher latitudes may have retained flowering patterns closer to the ancestral state, while rear-edge populations may have adapted to climates that have warmed over the last millennia such that they are now subtropical. Reduced vernalization requirements and shifts in phenology at the rear edge may thus be a relatively recent, post-glacial adaptation, diverging from an ancestral state of relying on winter cues to regulate phenology. Despite historical and contemporary climate change progressing at very different temporal scales, the evolution of reduced vernalization requirements since the LGM suggests that evolution of lower vernalization requirements in response to warming climate may be possible in populations at mid- and high latitude, and may be facilitated by genetic variation occurring at the rear-edge.

## Acknowledgements

This work was supported by the Swiss National Foundation (P2BSP3_195363) and the University of Virginia College of Arts and Sciences. We are grateful to David Westneat (University of Kentucky Ecological Research and Education Center Field Station, Lexington, KY), Gordon Burghardt (Blaine, TN), Trevor Stamey (Clemson University Experimental Forest, Clemson, SC), Jayne Lampley and John Walton (University of Alabama Tanglewood Station, Akron, AL), Theresa Jepsen (Florida State University Mission Road Research Facility, Tallahassee, FL) for logistical support in establishing field sites. For their help in raising of plants, collecting data, performing crosses and setting up the common garden experiments we thank A. Burricks, C. Claussen, S. Cox, L. Elhady, M. Gower-Fici, S. Kelly, O. Keenan, A. López, K. Lamb, H. Makowski, M. Marcich & E. Scott. We are also grateful to D. Brown, J. Collins, A. Diamond, L. Elliott, E. Galloway, F. Griffith, I. Guenther, J. Hansen, H. Horne, J. Kees, M. Kohout, R. Laporte, D. Reed and B. Sutherland for seed collection in natural populations. Collection permits were provided by the Florida Department of Environmental Protection, the Tennessee Department of Environment and Conservation, and Missouri Department of Conservation.

## Author Contributions

All authors contributed to the study design. AP collected seeds in the field and performed crosses, AP and MT raised plants and collected data, AP analyzed the data and wrote the manuscript with input from LG.

## Data availability statement

The data that support the findings of this study are openly available in Zenodo at http://doi.org/10.5281/zenodo.14773408.

## Supporting information

**Fig. S1:** Pruning of iNaturalist observations used in the Observational Study

**Fig. S2:** Example of reproductive stages of *Campanula americana* attributed to each observation

**Fig. S3:** Design of Experiment 3

**Fig. S4:** Competing models describing the variation in flowering time across latitudes

**Fig. S5:** Variation in time to bolting (a) and time between bolting and flowering (b) in Experiment 1

**Fig. S6:** Proportion of *Campanula americana* plants transitioning to bolting (a), from bolting to flowering (b) and the mean week of first flower (c) in Experiment 2

**Fig. S7:** Proportion of plants transitioning to bolting (a) and from bolting to flowering (b), and time to bolting (c) and between bolting and flowering (d) in Experiment 4

**Table S1** Estimation of first flower date in natural populations based on flowering stage

**Table S2a:** Description of the 17 climatic variables used in the Observational Study

**Table S2b:** Contribution to the discriminant function and range of the 17 climatic variables used in the Observational Study

**Table S3:** *Campanula americana* populations used in Experiments 1, 2 and 3

**Table S4a:** Dependent variables analyzed in Experiments 1 & 3

**Table S4b:** Dependent variables analyzed in Experiment 2

**Table S5:** Comparison of model fit for linear and quadratic relationship with latitude for Experiments 1, 2 and 3

**Table S6a:** Conditions in each common garden

**Table S6b:** Dates of transplanting and phenology estimation in each common garden

**Table S7:** Comparison of three models describing variation in the day of first flower across latitude

**Table S8:** Estimates of the effect of latitude, cohort and their interaction on traits in Experiment 1

**Table S9a:** Estimates of the effect of latitude on traits in Experiment 2

**Table S9b:** Comparison of traits between common gardens in Experiment 2

**Table S9c:** Comparison of the effect of latitude on traits between common gardens in Experiment 2

**Table S10a:** Estimates of the effect of latitude across three vernalization treatments on traits in Experiment 3

**Table S10b**: Comparison of the effect of vernalization treatments on traits in Experiment 4

**Table S10c:** Comparison of the effect of latitude on traits between vernalization treatments in Experiment 3

**Methods S1:** Selection of observations for estimation of day of first flower

**Methods S2:** Parametrization of analyses

**Methods S3:** Raising of *Campanula americana* plants and seed rearing in Experiment 1

**Methods S4:** Parametrization of the hierarchical mixed-effect models used in Experiments 1, 2, 3

**Methods S5:** Estimation of first flowering time of *Campanula americana* in common gardens

