## Supplementary for "Shifts in vernalization and phenology at the rear edge hold insight into adaptation of temperate plants to future milder winters"

Article acceptance date: 28 January 2025

The following Supporting Information is available for this article:

**Fig. S1:** Pruning of iNaturalist observations used in the Observational Study

**Fig. S2:** Example of reproductive stages of *Campanula americana* attributed to each observation

**Fig. S3:** Design of Experiment 3

**Fig. S4:** Competing models describing the variation in flowering time across latitudes

**Fig. S5:** Variation in time to bolting (a) and time between bolting and flowering (b) in Experiment 1

**Fig. S6:** Proportion of *Campanula americana* plants transitioning to bolting (a), from bolting to flowering (b) and the mean week of first flower (c) in Experiment 2

**Fig. S7:** Proportion of plants transitioning to bolting (a) and from bolting to flowering (b), and time to bolting (c) and between bolting and flowering (d) in Experiment 4

**Table S1** Estimation of first flower date in natural populations based on flowering stage

**Table S2a:** Description of the 17 climatic variables used in the Observational Study

**Table S2b:** Contribution to the discriminant function and range of the 17 climatic variables used in the Observational Study

**Table S3:** *Campanula americana* populations used in Experiments 1, 2 and 3

**Table S4a:** Dependent variables analyzed in Experiments 1 & 3

**Table S4b:** Dependent variables analyzed in Experiment 2

**Table S5:** Comparison of model fit for linear and quadratic relationship with latitude for Experiments 1, 2 and 3

**Table S6a:** Conditions in each common garden

**Table S6b:** Dates of transplanting and phenology estimation in each common garden

**Table S7:** Comparison of three models describing variation in the day of first flower across latitude

**Table S8:** Estimates of the effect of latitude, cohort and their interaction on traits in Experiment 1

**Table S9a:** Estimates of the effect of latitude on traits in Experiment 2

**Table S9b:** Comparison of traits between common gardens in Experiment 2

**Table S9c:** Comparison of the effect of latitude on traits between common gardens in Experiment 2

**Table S10a:** Estimates of the effect of latitude across three vernalization treatments on traits in Experiment 3

**Table S10b**: Comparison of the effect of vernalization treatments on traits in Experiment 4

**Table S10c:** Comparison of the effect of latitude on traits between vernalization treatments in Experiment 3

**Methods S1:** Selection of observations for estimation of day of first flower

**Methods S2:** Parametrization of analyses

**Methods S3:** Raising of *Campanula americana* plants and seed rearing in Experiment 1

**Methods S4:** Parametrization of the hierarchical mixed-effect models used in Experiments 1, 2, 3

**Methods S5:** Estimation of first flowering time of *Campanula americana* in common gardens

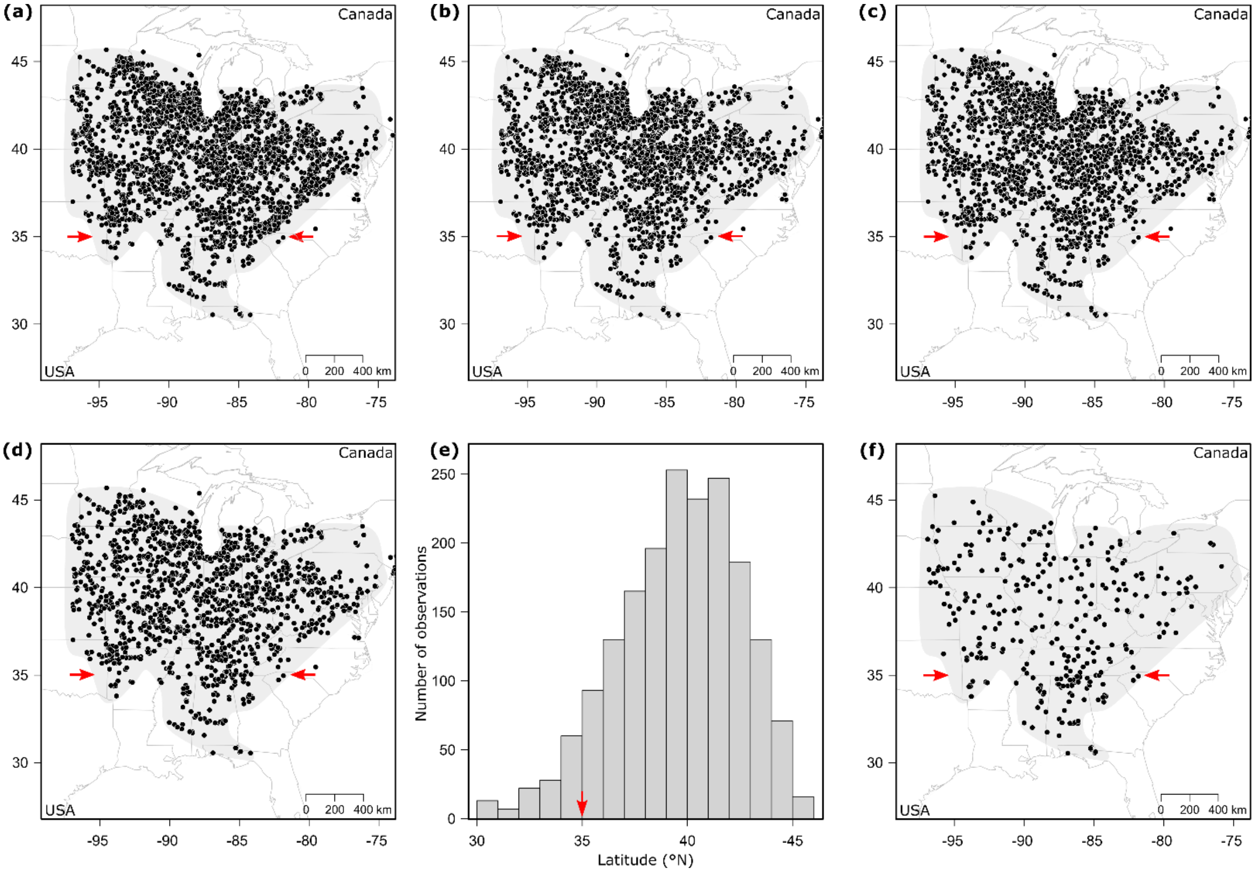


**Fig. S1: Pruning of iNaturalist observations used in the Observational Study.** The known range of natural populations of *Campanula americana* is indicated by grey shading; red arrows represent the latitudinal delimitation of the rear edge. Observations are indicated by black dots. (**a**) All observations. Observations remaining after removing high elevation observations and pruning to (**b**) 1/10 km grid/year, (**c**) 1/25 km grid/year or (**d**)1/50 km grid/year. (**e**) Density of observations across latitudes in (**d**). (**f**) Observations used to assess the day of first flower after pruning to get a more even distribution across latitude.

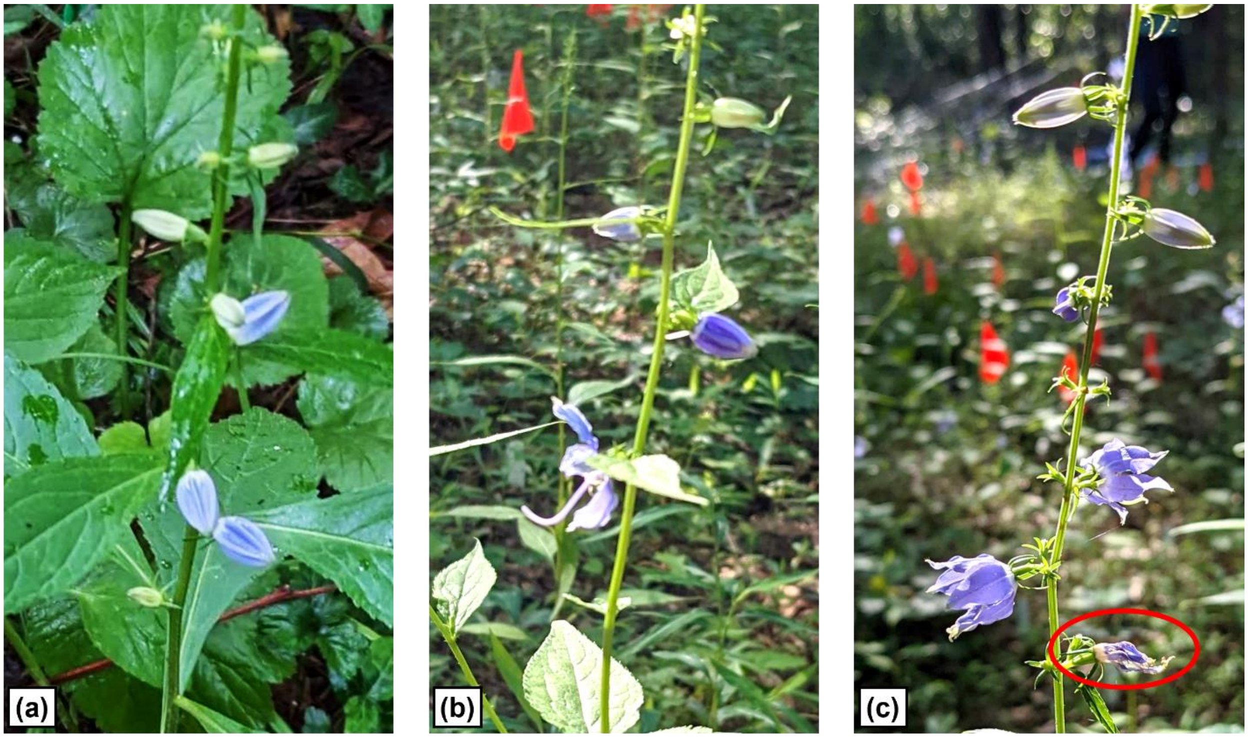


**Fig. S2: Examples of reproductive stage of *Campanula americana* attributed to observations.** **(a)** Stage 1, “one week before first flower”. **(b)** Stage 2, “week of first flower”. **(c)** Stage 3, “one week after first flower” with week-old dry flower circled in red. (Photographs by A. Perrier in 2023 from field experiments.)

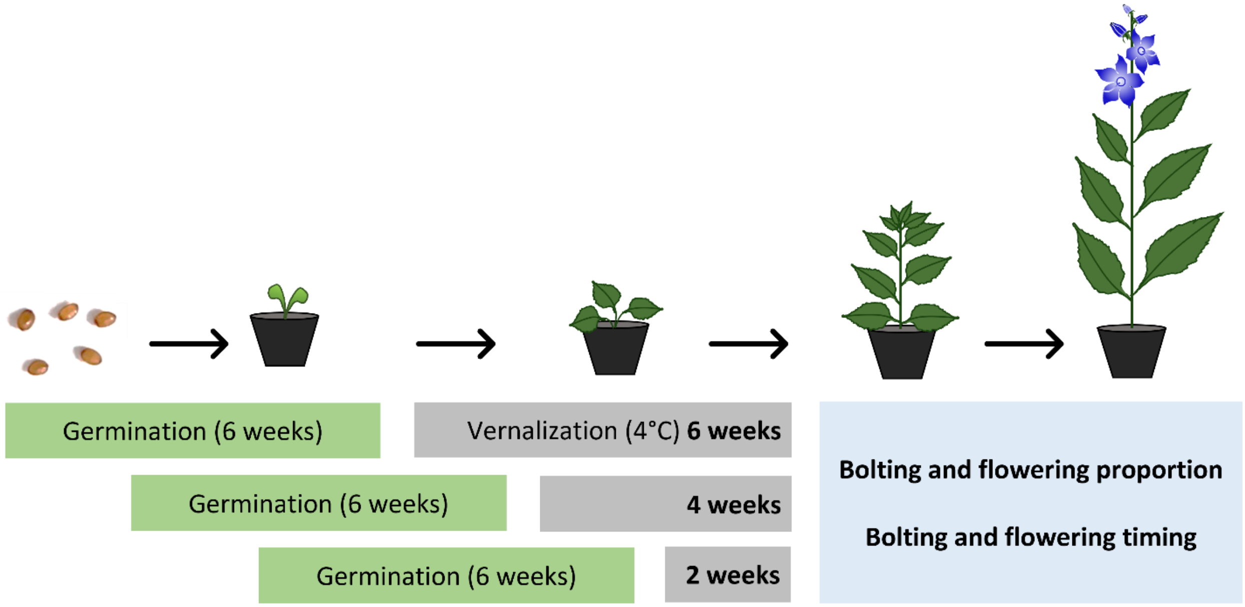


**Fig. S3: Design of Experiment 3.** Seedlings of 12 populations from across the range of *Campanula americana* were raised in growth chambers (germination and vernalization) and the greenhouse (bolting and flowering) and exposed to one of three vernalization treatments (six, four or two weeks, 540 plants per treatment, 1620 total). Sowing was timed such that all treatments ended vernalization at the same time. The proportion and timing of bolting and flowering was recorded three times a week after vernalization.

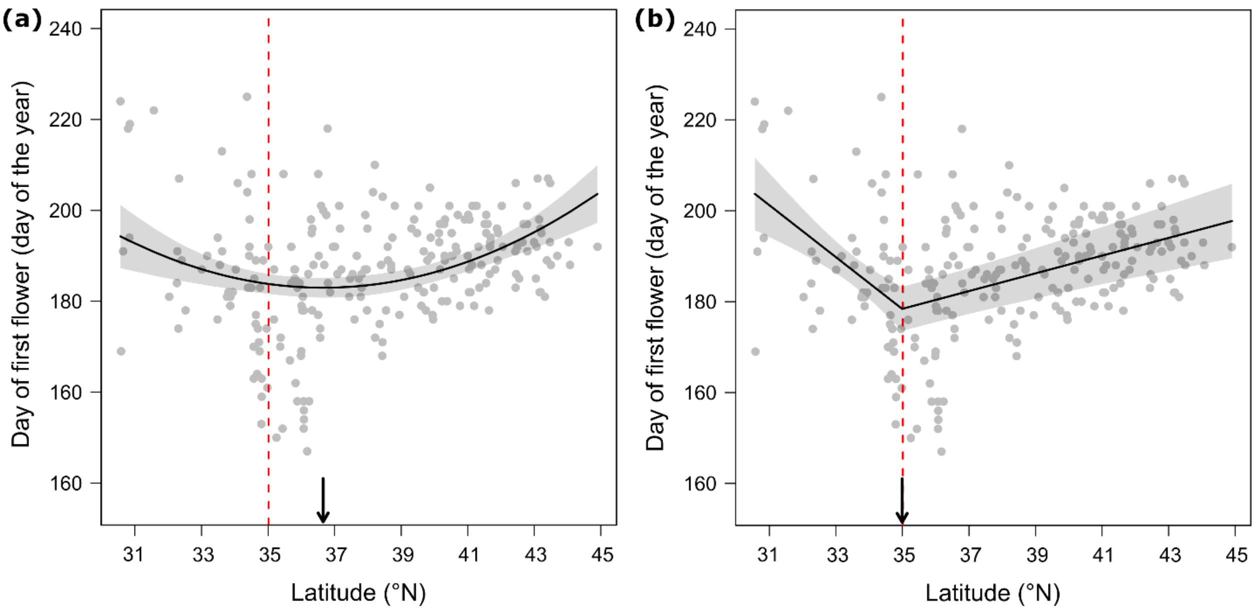


**Fig. S4: Competing models describing the variation in flowering time across latitudes.** The day of first flower was inferred based on iNaturalist observations recorded across the range of *Campanula americana* between 2018 and 2022. The red dashed lines represent the *a priori* delimitation of the rear edge (35 °N). Variation in the day of first flower was best described either by a quadratic model (**a**) or a piecewise model (**b**). Solid lines represent the significant model-predicted slope of the relationship with latitude, with shading indicating the 95% confidence. Arrows indicate the latitude of the earliest day of first flower predicted by the polynomial model (**a**) or the breakpoint identified by the piecewise model (**b**). Test statistics in Table S7.

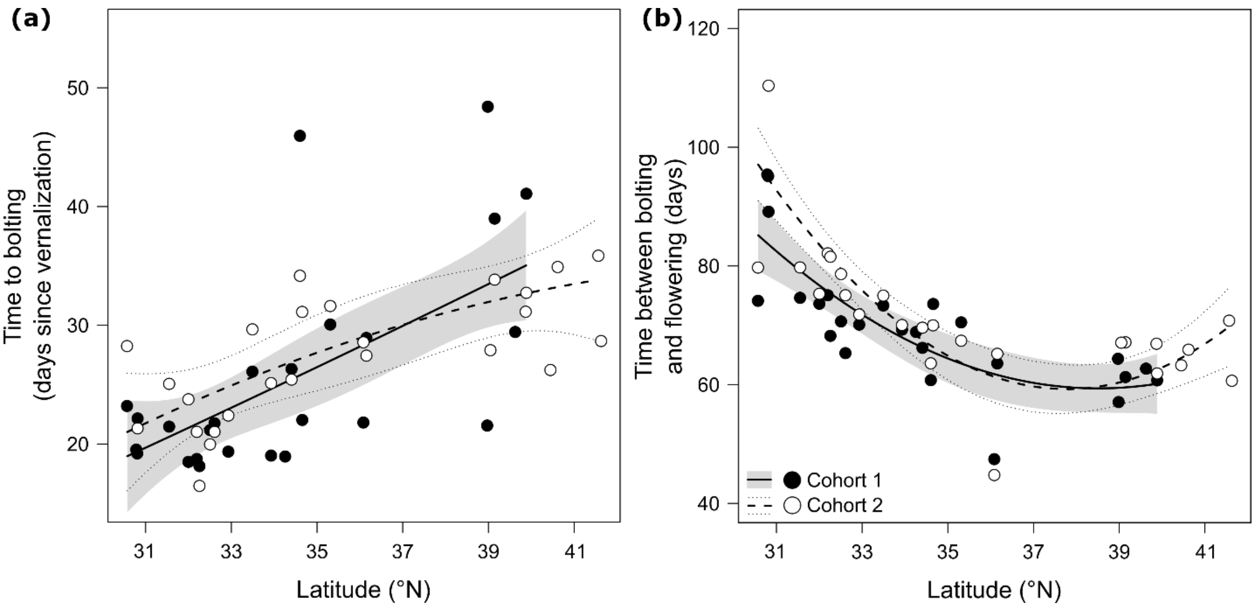


**Fig. S5: Variation in time to bolting (a) and time between bolting and flowering (b) in Experiment 1.** Both traits were recorded for populations sampled across the range of *Campanula americana* and raised in a common controlled environment in two cohorts. Lines represent the significant model-predicted relationship between each trait estimated at the individual level and latitude for each cohort, with the 95% confidence interval indicated as shading (Cohort 1) or dotted lines (Cohort 2). Dots represent the trait mean of each population. Test statistics are reported in Table 2 and Table S8.

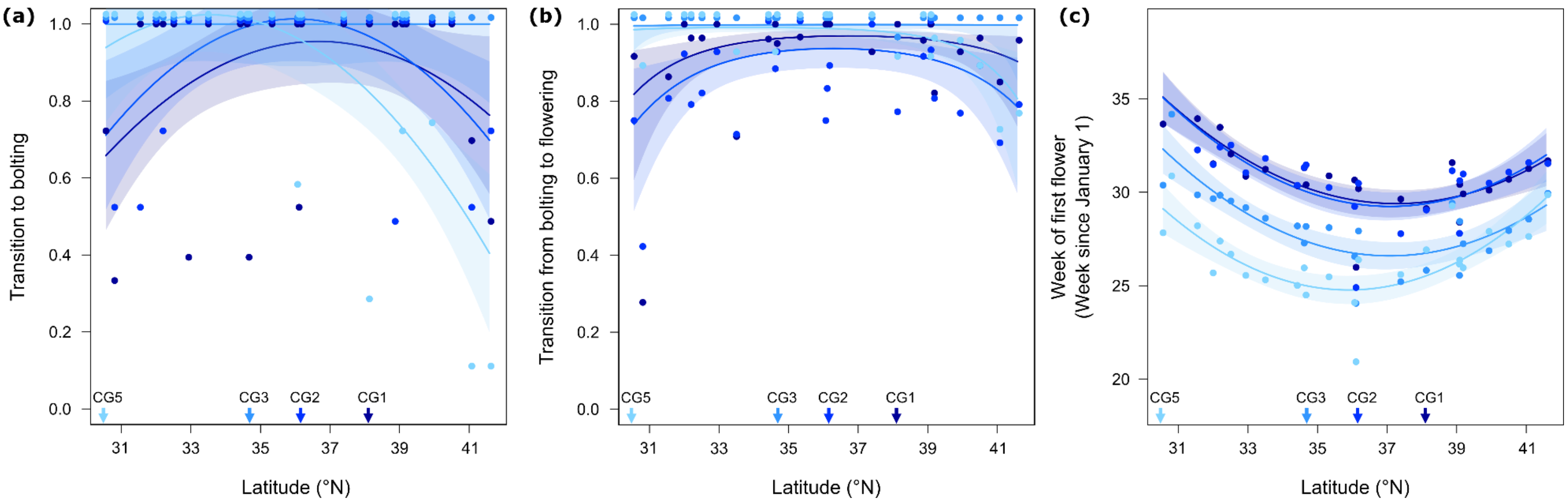


**Fig. S6: Proportion of *Campanula americana* plants transitioning to bolting (a), from bolting to flowering (b) and the mean week of first flower (c) in Experiment 2.** Colors distinguish the gardens, and their latitude is indicated with arrows along the x-axis. Lines represent the significant model-predicted relationship between each trait estimated at the individual level (population level for transition to bolting) and latitude in each garden, with the 95% confidence interval indicated as shading. Dots represent the trait mean of each population. In (a) and (b), trait means equal to 1 were offset for clarity, resulting in some points above 1.0. Test statistics are reported in Table 2 and Tables S9.

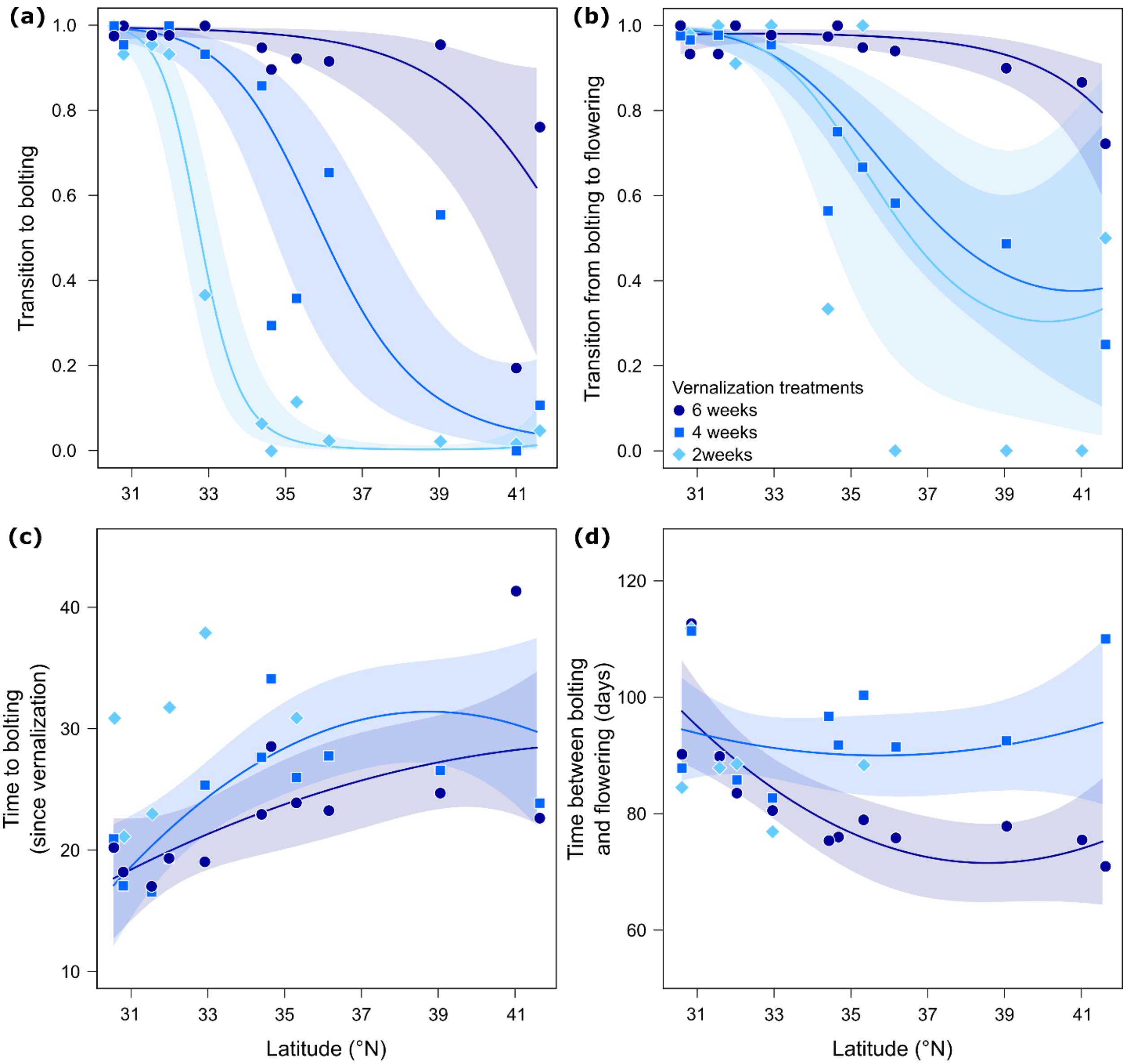


**Fig. S7: Proportion of plants transitioning to bolting (a) and from bolting to flowering (b), and time to bolting (c) and between bolting and flowering (d) in Experiment 3**. Populations sampled across the range of *Campanula americana* and exposed to six, four or two week vernalization treatments in controlled conditions. For traits related to timing, the 2-week treatment was excluded from analysis as too few individuals bolted successfully (population means based on at least five individuals shown for comparison). Lines represent the significant model-predicted relationship for each trait estimated at the individual level and population latitude in each vernalization treatment; 95% confidence interval indicated as shading. Dots represent the trait means of each population. Test statistics are reported in Table 2 and Tables S10.

**Table S1: Estimation of first flower date in natural populations based on reproductive stage**

| Stage | Criteria | First flower date = Observation date |
|---|---|---|
| 1 – One week before first flower | Large purple buds, no open flowers (Fig. S2a) | +7 |
| 2 – Week of first flower | First flowers open, no dried flowers (Fig. S2b) | 0 |
| 3 – One week after first flower | First flowers dry but petals still colored, no fruits (Fig. S2c) | -7 |

**Table S2a: Description of the 17 climatic variables used in the Observational Study**

| Climatic variable | Abbreviation | Description |
|---|---|---|
| Length of vernalization | LVerna | Number of days of cold, *Tmax* < 4° C (September 1st of the prior year to date of observation) |
| Temperature in spring | TSpring | Average *Tmean* in March, April and May during the year of observation |
| Temperature in summer | TSummer | Average *Tmean* in June, July and August during the year of observation |
| Temperature before flowering | TBFlow | Average *Tmean* during the 30 days before first flower |
| Temperature after flowering | TAFlow | Average *Tmean* during the 30 days after first flower |
| Precipitation before flowering | PBFlow | Sum of precipitation 30 days before first flower |
| Precipitation after flowering | PAFlow | Sum of precipitation 30 days after first flower |
| Warmest day of the year | DWarm | Day of the year of observation with highest *Tmax* |
| Wettest 10 days of summer | DWet | Fifth day of the 10 consecutive days with highest sum of precipitation during the warmest half of the year (April 1st to September 30th) |
| Driest 10 days of summer | DDry | Fifth day of the 10 consecutive days with lowest sum of precipitation during the warmest half of the year (April 1st to September 30th) |
| Time between flowering and | | |
| -the end of vernalization | FlowV | Number of days between first flower and the last day where *Tmax* < 4° C * |
| -the start of the growing season | FlowGS | Number of days between first flower and the last day where *Tmean* < 10° C |
| -the warmest day of the year | FlowWarm | Number of days between first flower and the warmest day of the year (DWarm) |
| -the wettest day of the year | FlowWet | Number of days between first flower and the wettest 10 days of summer (DWet) |
| -the driest day of the year | FlowDry | Number of days between first flower and the driest 10 days of summer (DDry) |
| Day length at flowering | DayLength | Day length (in hours) on the day of first flower |
| Day length at the start of the growing season | DayLengthGS | Day length (in hours) on the last day where *Tmean* < 10° C |

For each observation, we obtained mean, maximum, and minimum daily temperature (*Tmean*, *Tmax*, *Tmin*) and daily precipitation (*Prec*) for the day of the year of observation and the year before, recorded at the observations coordinates from the PRISM database (https://prism.oregonstate.edu, accessed 01/09/2023) using the *prism* R package (Edmund & Bell, 2015).

* For eight populations, *Tmax* never dropped below 4° C in winter (*i.e.* LVerna = 0), and FlowV was calculated relative to the coldest day in the winter.

**Table S2b: Contribution to the discriminant function and range of the 17 climatic variables used in the Observational Study**

| Climatic variable | Unit | Min. | Mean | Max. | Contribution |
|---|---|---|---|---|---|
| LVerna | Days | 0 | 42 | 121 | 0.62834 |
| FlowGS | Days | 16 | 63 | 158 | 0.30282 |
| FlowV | Days | 54 | 115.3 | 226 | 0.04068 |
| PBFlow | mm | 24 | 126.9 | 409.6 | 0.00828 |
| TSpring | ° C | 5.7 | 12.6 | 21.1 | 0.00643 |
| FlowWarm | Days | -112 | -17 | 52 | 0.00252 |
| TAFlow | ° C | 20.9 | 24.7 | 29.7 | 0.00200 |
| TSummer | ° C | 20.6 | 24.2 | 28.1 | 0.00187 |
| DDry | Days | 91 | 186 | 271 | 0.00160 |
| PAFlow | mm | 5.3 | 110 | 554.7 | 0.00122 |
| DWarm | Days | 147 | 204 | 264 | 0.00118 |
| TBFlow | ° C | 16.7 | 23.6 | 29.2 | 0.00113 |
| FlowWet | Days | -114 | 3 | 107 | 0.00064 |
| FlowDry | Days | -102 | 1 | 118 | 0.00058 |
| DayLengthGS | Hours | 11.5 | 13.9 | 15.2 | 0.00057 |
| DWet | Days | 91 | 184 | 274 | 0.00009 |
| DayLength | Hours | 13.3 | 14.7 | 15.4 | 0.00008 |

**Table S3: *Campanula americana* populations used in Experiments 1, 2 and 3**

| | | | Experiment 1 | | | | Experiment 2 (per garden) | | | Experiment 3 (per treatment) | |
|---|---|---|---|---|---|---|---|---|---|---|---|
| | Latitude | Longitude | Cohort 1 | | Cohort 2 | | | | | | |
| ID | [° N] | [° E] | Families | Individuals | Families | Individuals | Seed type | Families | Individuals | Families | Individuals |
| FL83 | 30.56468 | -84.95982 | 20 | 21 | 14 | 20 | Crosses | 15 | 30 | 14 | 45 |
| FL2 | 30.78036 | -85.16838 | 12 | 18 | - | - | - | - | - | - | - |
| FL1 | 30.80288 | -85.24966 | 16 | 21 | - | - | - | - | - | - | - |
| FL81 | 30.81165 | -85.22535 | 18 | 18 | 13 | 19 | Crosses | 15 | 30 | 15 | 45 |
| AL2 | 31.54793 | -87.51453 | 20 | 23 | 15 | 25 | Crosses | 15 | 30 | 15 | 45 |
| MS6 | 31.99803 | -89.35603 | 20 | 23 | 14 | 25 | Crosses | 15 | 30 | 15 | 45 |
| AL23 | 32.19395 | -86.78484 | 16 | 21 | 14 | 23 | Crosses | 15 | 30 | - | - |
| AL8 | 32.25610 | -86.49330 | 20 | 20 | 15 | 20 | - | - | - | - | - |
| AL22 | 32.50376 | -87.50489 | 19 | 29 | 8 | 20 | Crosses | 15 | 30 | - | - |
| AL19 | 32.60700 | -88.19192 | 20 | 28 | 12 | 20 | - | - | - | - | - |
| AL79 | 32.92932 | -88.20820 | 19 | 26 | 13 | 24 | Crosses | 15 | 30 | 15 | 45 |
| AL21 | 33.49109 | -86.79829 | 20 | 27 | 13 | 20 | Crosses | 15 | 30 | - | - |
| MS9 | 33.92778 | -89.01710 | 19 | 25 | 10 | 20 | - | - | - | - | - |
| MS1 | 34.25635 | -88.88169 | 19 | 23 | - | - | - | - | - | - | - |
| MS8 | 34.40427 | -88.83093 | 20 | 30 | 9 | 20 | Crosses | 15 | 30 | 13 | 45 |
| GA1 | 34.60074 | -84.69664 | 19 | 30 | 10 | 29 | Crosses | 15 | 30 | - | - |
| ALBG | 34.65346 | -86.51638 | 16 | 22 | 13 | 23 | Crosses | 15 | 30 | 13 | 45 |
| TN3 | 35.30932 | -90.06770 | 15 | 21 | 10 | 26 | Crosses | 15 | 30 | 13 | 45 |
| AR3 | 36.04148 | -92.06613 | - | - | | | Field | 11 | 30 | - | - |
| TN34 | 36.08222 | -86.29611 | 15 | 17 | 12 | 24 | Crosses | 15 | 30 | - | - |
| AR2 | 36.15235 | -94.30465 | 16 | 19 | 15 | 25 | Crosses | 15 | 30 | 14 | 45 |
| KY5 | 37.36005 | -84.77186 | - | - | - | - | Field | 15 | 30 | - | - |
| KY1 | 38.09215 | -84.98688 | - | - | - | - | Field | 15 | 30 | - | - |
| MO2 | 38.83025 | -92.28509 | - | - | - | - | Field | 15 | 30 | - | - |
| OH1 | 39.03935 | -84.32774 | - | - | - | - | Field | 15 | 30 | - | - |
| KS60 | 39.04742 | -95.68152 | 20 | 23 | 14 | 25 | Crosses | 15 | 30 | 15 | 45 |
| IN5 | 39.14583 | -86.54833 | 19 | 30 | 13 | 25 | Crosses | 15 | 30 | - | - |
| IN7 | 39.87011 | -86.16020 | - | - | 9 | 20 | - | - | - | - | - |
| OH119 | 39.88500 | -83.99700 | 18 | 29 | 8 | 20 | Crosses | 15 | 30 | - | - |
| IN4 | 40.44806 | -86.93444 | - | - | 11 | 20 | Crosses | 15 | 30 | - | - |
| IL10 | 40.61750 | -89.01778 | - | - | 12 | 20 | - | - | - | - | - |
| IN2 | 41.01731 | -85.23872 | - | - | - | - | Crosses | 15 | 30 | 10 | 45 |
| IA19 | 41.56611 | -90.47500 | - | - | 13 | 20 | Crosses | 15 | 30 | - | - |
| IN1 | 41.62995 | -85.90436 | - | - | 13 | 20 | - | - | - | - | - |

**Table S4a: Dependent variables analyzed in Experiments 1 & 3**

| Dependent variable | Distribution | Description |
|---|---|---|
| Flowering success | Binary | Production of at least one flower |
| -Transition to bolting * | Binary | Production of a flowering stalk |
| -Transition from bolting to flowering * | Binary | Production of at least one flower on a plant that bolted |
| Time to flowering | Gaussian | Number of days between the end of vernalization and opening of the first flower |
| -Time to bolting | Gaussian | Number of days between the end of vernalization and the day bolting was observed |
| -Time between bolting and flowering | Gaussian | Number of days between bolting and opening of the first flower |

* Only for Exp. 3.

**Table S4b: Dependent variables analyzed in Experiment 2**

| Dependent variable | Distribution | Description |
|---|---|---|
| Early flowering | Binary | Production of at least one bud by the first collection visit |
| -Transition to bolting † | Binary | Production of a flowering stalk |
| -Transition from bolting to flowering | Binary | Production of at least one bud on plants that bolted |
| Week of first flowering | Gaussian | Number of weeks since January $1^{st}$ 2023 until the first flower |
| Time to flowering | Gaussian | Number of weeks from transplant until the first flower |

Phenotypic data was collected over three successive collection rounds (only two for CG5). Success and timing estimates recorded on plants that died after that data collection round were still included in the analysis.

† Analyses at the individual level except *transition to bolting* for which population means within sites were analyzed, as some sites did not have enough variance for individual analysis.

**Table S5: Comparison of model fit for linear and quadratic relationship with latitude for Experiments 1, 2 and 3**

| Dependent variable | Effect of latitude | Fixed effect structure | *AICc* |
|---|---|---|---|
| *Experiment 1* | | | |
| Flowering success | **Linear** | **Latitude + cohort + (latitude *cohort)** | **581.03** |
| | Quadratic | Latitude + latitude$^2$ + cohort + (latitude + latitude$^2$ *cohort) | 585.03 |
| Time to flowering | Linear | Latitude + cohort + (latitude *cohort) | 6396.81 |
| | **Quadratic** | **Latitude + latitude$^2$ + cohort + (latitude + latitude$^2$ *cohort)** | **6343.74** |
| Time to bolting | Linear | Latitude + cohort + (latitude *cohort) | 6702.45 |
| | **Quadratic** | **Latitude + latitude$^2$ + cohort + (latitude + latitude$^2$ *cohort)** | **6669.84** |
| Time between bolting and flowering | Linear | Latitude + cohort + (latitude *cohort) | 6320.49 |
| | **Quadratic** | **Latitude + latitude$^2$ + cohort + (latitude + latitude$^2$ *cohort)** | **6256.20** |
| *Experiment 2* | | | |
| Early flowering | Linear | Latitude + common garden + (latitude *common garden) | 1716.92 |
| | **Quadratic** | **Latitude + latitude$^2$ + common garden + (latitude + latitude$^2$ *common garden)** | **1696.75** |
| Transition to Bolting | Linear | Latitude + common garden + (latitude *common garden) | -411.14 |
| | **Quadratic** | **Latitude + latitude$^2$ + common garden + (latitude + latitude$^2$ *common garden)** | **-419.91** |
| Transition from bolting to flowering | Linear | Latitude + common garden + (latitude *common garden) | 869.28 |
| | Quadratic | Latitude + latitude$^2$ + common garden + (latitude + latitude$^2$ *common garden) | 870.42 |
| Week of first flower | Linear | Latitude + common garden + (latitude *common garden) | 8926.99 |
| | **Quadratic** | **Latitude + latitude$^2$ + common garden + (latitude + latitude$^2$ *common garden)** | **8845.18** |
| Time to flowering (since transplanting) | Linear | Latitude + common garden + (latitude *common garden) | 8926.99 |
| | **Quadratic** | **Latitude + latitude$^2$ + common garden + (latitude + latitude$^2$ *common garden)** | **8843.18** |
| *Experiment 3* | | | |
| Flowering success | Linear | Latitude + treatment + (latitude *treatment) | 942.67 |
| | **Quadratic** | **Latitude + latitude$^2$ + treatment + (latitude + latitude$^2$ *treatment)** | **922.95** |
| Transition to Bolting | Linear | Latitude + treatment + (latitude *treatment) | 866.58 |
| | **Quadratic** | **Latitude + latitude$^2$ + treatment + (latitude + latitude$^2$ *treatment)** | **840.19** |
| Transition from bolting to flowering | Linear | Latitude + treatment + (latitude *treatment) | 458.84 |
| | **Quadratic** | **Latitude + latitude$^2$ + treatment + (latitude + latitude$^2$ *treatment)** | **455.55** |
| Time to flowering | Linear | Latitude + treatment + (latitude *treatment) | 5307.30 |
| | **Quadratic** | **Latitude + latitude$^2$ + treatment + (latitude + latitude$^2$ *treatment)** | **5259.39** |
| Time to bolting | Linear | Latitude + treatment + (latitude *treatment) | 5450.59 |
| | **Quadratic** | **Latitude + latitude$^2$ + treatment + (latitude + latitude$^2$ *treatment)** | **5410.99** |
| Time between bolting and flowering | Linear | Latitude + treatment + (latitude *treatment) | 5184.25 |
| | **Quadratic** | **Latitude + latitude$^2$ + treatment + (latitude + latitude$^2$ *treatment)** | **5139.68** |

The best model (bold) was identified by the lowest AICc values.

**Table S6a: Conditions in each common garden**

| | Latitude (° N) | Longitude (° E) | US state | Days of vernalization (< 4 °C*) | Minimum temperature in spring** | Maximum temperature in summer** |
|---|---|---|---|---|---|---|
| CG1 | 38.08079 | -84.47130 | KY | 7 | 10.33 | 26.77 |
| CG2 | 36.18525 | -83.69820 | TN | 7 | 9.21 | 26.77 |
| CG3 | 34.68746 | -82.87249 | SC | 4 | 10.77 | 28.67 |
| CG5 | 30.45723 | -84.33320 | FL | 3 | 13.91 | 30.60 |

* Based on daily minimum temperature recorded in each site from transplant until the end of the experiment
** Average of daily minimum or maximum temperature recorded before (spring) or after (summer) first flower in each site.

**Table S6b: Dates of transplanting and visits for estimation of first flowering time for each common garden**

| | Transplanting | Visit 1 | Visit 2 | Visit 3 |
|---|---|---|---|---|
| CG1 | 08/04/2023 | 16/07/2023 | 12/08/2023 | 09/09/2023 |
| CG2 | 05/04/2023 | 15/07/2023 | 18/08/2023 | 08/09/2023 |
| CG3 | 22/03/2023 | 14/07/2023 | 13/08/2023 | 06/09/2023 |
| CG5 | 02/03/2023 | 12/07/2023 | 17/08/2023 | - |

**Table S7: Comparison of three models describing variation in the day of first flower across latitude**

| Model | *P* | *AICc* | Minimum (°N) |
|---|---|---|---|
| Piecewise | <0.001 | 1879.84 | 34.98 |
| Polynomial | <0.001 | 1880.18 | 36.64 |
| Linear | 0.002 | 1911.96 | - |

The dependent variable was the day of first flower inferred from observations. All three models tested for an effect of latitude, but each model integrated this effect differently (see Supporting method S2a). The minimum represents the latitude of the earliest predicted day of first flower.

**Table S8: Estimates of the effect of latitude, cohort and their interaction on traits in Experiment 1**

| | Latitude (L) | | | | Cohort (C) | | L * C | |
|---|---|---|---|---|---|---|---|---|
| Dependent variable | *β C1* | *β C2* | *β² C1* | *β² C2* | *C1 vs C2* | | *β C1 vs C2* | |
| Flowering success | -0.02 | -0.01 | - | - | **-3.50** | *** | 0.23 | (*) |
| Time to flowering | -32.53 | -49.71 | 0.45 | 0.67 | **-2.94** | **, * | **1.47** | *** |
| Time to bolting | 1.26 | 4.92 | 0.01 | -0.05 | -1.21 | | 0.45 | |
| Time between bolting and flowering | -31.53 | -55.10 | 0.41 | 0.73 | -0.50 | | **1.19** | *** |

Values depict the estimated slope coefficients of the linear (*β*) and quadratic (*β²*) effect of latitude for each cohort (C), the estimated pairwise difference between cohorts and the estimated pairwise contrast of the effect of latitude between cohorts (based on the latitude – cohort interaction). The latter two estimates were obtained from Tukey's tests (*P*-values < 0.05 bolded); (*) *P*<0.1, * *P*<0.05, ** *P*<0.01, *** *P*<0.001. A Bonferroni correction was applied to the significance of each fixed effect for traits that comprise time to flowering as they are non-independent. Any changes in significance thresholds are indicated as a secondary *P*-value.

**Table S9a: Estimates of the effect of latitude on traits in Experiment 2**

| | Latitude | | | | Latitude$^2$ | | | |
|---|---|---|---|---|---|---|---|---|
| Dependent variable | *CG1* | *CG2* | *CG3* | *CG5* | *CG1* | *CG2* | *CG3* | *CG5* |
| Early flowering | 1.08 | 1.27 | 0.44 | 0.38 | -0.01 | -0.02 | -0.01 | -0.01 |
| Transition to bolting | 0.08 | 0.11 | 0.00 | 0.10 | -1e-3 | -1e-3 | 0.00 | -1e-3 |
| Transition from bolting to flowering | 0.22 | 0.37 | 0.01 | 0.12 | -3e-3 | -5e-3 | -8e-5 | -2e-3 |
| Week of first flower | -9.51 | -10.23 | -10.03 | -11.06 | 0.13 | 0.14 | 0.14 | 0.15 |
| Time to flowering (since transplanting) | -9.51 | -10.23 | -10.03 | -11.06 | 0.13 | 0.14 | 0.14 | 0.15 |

Values depict the estimated slope coefficients of the linear and quadratic effect of latitude in each common garden (CG).

**Table S9b: Comparison of traits between common gardens in Experiment 2**

| | Common garden | | | | | |
|---|---|---|---|---|---|---|
| Dependent variable | *1 vs 2* | *1 vs 3* | *1 vs 5* | *2 vs 3* | *2 vs 5* | *3 vs 5* |
| Early flowering | 0.23 | **-2.91** *** | **-1.21** *** | **-3.14** *** | **-1.43** *** | **1.71** ** |
| Transition to bolting | -0.01 | -0.01 | -0.00 | 0.00 | 0.01 | 0.00 |
| Transition from bolting to flowering | 0.74 | **-3.40** * | -1.13 | **-4.14** ** | **-1.87** ***, ** | 2.27 |
| Week of first flower | 0.19 | **2.84** *** | **4.84** *** | **2.65** *** | **4.65** *** | **2.00** *** |
| Time to flowering (since transplanting) | -0.23 | 0.41 | **-0.59** * | **0.65** **, * | -0.35 | **-1.00** *** |

Values depict the estimated pairwise differences between common gardens obtained from Tukey's tests. Estimates with *P*-values < 0.05 in bold; * *P*<0.05, ** *P*<0.01, *** *P*<0.001. A Bonferroni correction was applied to the significance of each fixed effect for traits that comprise Early flowering and for the two timing traits, as they are non-independent. Any changes in significance thresholds are indicated as a secondary *P*-value.

**Table S9c: Comparison of the effect of latitude on traits between common gardens in Experiment 2**

| | Latitude * common garden | | | | | |
|---|---|---|---|---|---|---|
| Dependent variable | *1 vs 2* | *1 vs 3* | *1 vs 5* | *2 vs 3* | *2 vs 5* | *3 vs 5* |
| Early flowering | -0.03 | -0.00 | **0.41** *** | 0.03 | **0.44** *** | **0.41** *** |
| Transition to bolting | 0.00 | 0.00 | **0.01** ** | 0.00 | **0.01** * | **0.01** *, (*) |
| Transition from bolting to flowering | 0.04 | -0.00 | **0.32** ***, ** | -0.04 | **0.23** ** | 0.33 |
| Week of first flower | -0.02 | -0.03 | **-0.35** *** | -0.01 | **-0.33** *** | **-0.32** *** |
| Time to flowering (since transplanting) | -0.02 | -0.03 | **-0.35** *** | -0.01 | **-0.33** *** | **-0.32** *** |

Values depict the estimated pairwise contrast of the effect of latitude between common gardens (based on the latitude–garden interaction) obtained from Tukey's tests. Estimates with *P*-values < 0.05 in bold; * *P*<0.05, ** *P*<0.01, *** *P*<0.001. A Bonferroni correction was applied to the significance of each fixed effect for traits that comprise Early flowering and for the two timing traits, as they are non-independent. Any changes in significance thresholds are indicated as a secondary *P*-value.

**Table S10a: Estimates of the effect of latitude across three vernalization treatments on traits in Experiment 3**

| | Latitude | | | Latitude$^2$ | | |
|---|---|---|---|---|---|---|
| Dependent variable | *V6* | *V4* | *V2* | *V6* | *V4* | *V2* |
| Flowering success | 0.12 | -0.31 | -0.53 | -0.00 | 0.00 | 0.01 |
| Transition to bolting | 0.04 | -0.26 | -0.58 | -0.00 | 0.00 | 0.01 |
| Transition from bolting to flowering | 0.08 | -0.35 | -0.15 | -0.00 | 0.00 | 0.00 |
| Time to flowering | -25.95 | 1.11 | - | 0.35 | 0.00 | - |
| Time to bolting | 5.30 | 16.47 | - | -0.06 | -0.21 | - |
| Time between bolting and flowering | -31.48 | -11.96 | - | 0.41 | 0.17 | - |

Values depict the estimated slope coefficients of the linear and quadratic effect of latitude in each vernalization treatment (V). For traits related to timing, the 2-week vernalization treatment was removed from analysis as too few individuals bolted successfully.

**Table S10b: Comparison of the effect of vernalization treatments on traits in Experiment 3**

| | Treatment | | | | | |
|---|---|---|---|---|---|---|
| Dependent variable | *V6 vs V4* | | *V6 vs V2* | | *V4 vs V2* | |
| Flowering success | **3.04** | *** | **6.54** | *** | **3.49** | *** |
| Transition to bolting | **2.95** | *** | **7.17** | *** | **4.22** | *** |
| Transition from bolting to flowering | **1.74** | *** | **1.85** | ** | 0.11 | |
| Time to flowering | **-12.10** | *** | - | | - | |
| Time to bolting | **-3.91** | *** | - | | - | |
| Time between bolting and flowering | **-9.35** | *** | - | | - | |

Values depict the estimated pairwise differences between treatments obtained from Tukey's tests. Estimates with *P*-values < 0.05 in bold; ** *P*<0.01, *** *P*<0.001. A Bonferroni correction was applied to the significance of each fixed effect for traits that comprise flowering success and time to flowering as they are non-independent, but significance thresholds did not change. For traits related to timing, the 2-week treatment was removed from analysis as too few individuals bolted successfully.

**Table S10c: Comparison of the effect of latitude on traits between vernalization treatments in Experiment 3**

| | Latitude * Treatment | | | | | |
|---|---|---|---|---|---|---|
| Dependent variable | *V6 vs V4* | | *V6 vs V2* | | *V4 vs V2* | |
| Flowering success | **0.52** | ** | **0.89** | *** | **0.37** | ** |
| Transition to bolting | **0.42** | **, * | **0.78** | *** | **0.36** | **, * |
| Transition from bolting to flowering | **0.67** | ** | **0.82** | **, * | 0.14 | |
| Time to flowering | **-4.10** | *** | - | | - | |
| Time to bolting | **-0.82** | *** | - | | - | |
| Time between bolting and flowering | **-3.31** | *** | - | | - | |

Values depict the estimated pairwise contrast of the effect of latitude between treatments (based on the latitude – treatment interaction) obtained from Tukey's tests. Estimates with *P*-values < 0.05 in bold; * *P*<0.05, ** *P*<0.01, *** *P*<0.001. A Bonferroni correction was applied to the significance of each fixed effect for traits that comprise flowering success and time to flowering as they are non-independent. Any changes in significance thresholds are indicated as a secondary *P*-value. For traits related to timing, the 2-week treatment was removed from analysis as too few individuals bolted successfully.

**Methods S1: Selection of observations for estimation of day of first flower**

To develop a set of observations from which to estimate the day of first flower, we first created a list of all observations of *Campanula americana* in iNaturalist (https://www.inaturalist.org) recorded between 2018 and 2022 using the built in export tool (https://www.inaturalist.org/observations/export, accessed 23/02/2023). Export was set to the following criteria: observation with photograph, "research" quality grade, "most agree" with identification, and not cultivated, resulting in a set of 10,250 observations (Fig. S1a). We then removed observations that occurred at elevations higher than 600m to avoid the confounding effect of similar environmental gradients that occur with latitude and elevation (9697 observations retained, ~95%). This also removed populations belonging to two smaller genetic lineages that occur in the Appalachian Mountains and that do not occupy southern glacial refugia (Barnard-Kubow *et al.*, 2015).

Observations gathered from publicly available data such as iNaturalist are often not sampled equally across the range. To address unequal sampling, we first pruned observations to limit the bias of denser sampling closer to cities and to remove potential duplicates within a population. Pruning was accomplished by creating a 50 by 50 km grid across the whole range and randomly selecting one observation per grid cell for each year of observation to retain homogeneous coverage across the range in each year (Fig. S1d; 1849 observations retained). The size of the grid was determined by testing pruning with increasing grid sizes (10km, 25km, 50km) until no spatial clustering, e.g. around cities, was visually observed (Fig S1b, S1c). Further pruning included removing observations outside of the known natural range (attributed to errors in the observations' coordinates or cultivated individuals), as well as observations recorded outside of the known flowering period (May 1 to September 1, personal obs.). Finally, there was a strong numerical bias towards high latitude in this dataset because the species' range is smaller and populations are less dense at low latitudes (Fig. S1e). Because we use latitude as main explanatory variable in this study, we adjusted for this bias by retaining all observations in the lower latitudinal third, i.e. below 35°N, and randomly selecting twice that number of populations in the upper two-thirds of the distribution within each year. This effectively homogenized observations across latitudes and resulted in a final sample of 335 observations for which flowering phenology was estimated (Fig. S1f).

To infer the day of first flower, we first characterized the reproductive stage of each iNaturalist observation by manually inspecting the associated photographs (Table S1, Fig, S2). We only kept observations for which assessing the reproductive stage was possible (i.e. clear photograph with flowers or buds visible). We inferred reproductive stages on observations where plants were close to the initiation of flowering (one week before or after first flower, Table S1), as these are easiest to assess from photographs (Fig, S2). We discarded observations that were clearly not in a natural environment based on photographs (urban park, private garden). This resulted in a final dataset of 238 observations for analysis (Fig 1a). The day of first flower was inferred for each observation by adjusting the observation date by the estimated number of days between observation and the estimated time of first flower based on reproductive stages (Table S1).

**Methods S2: Parametrization of analyses**

**S2a: Parametrization of the three hierarchical mixed-effect models testing the relationship between the day of first flower and latitude.**

The first two models were parametrized using the R packages *lme4* (Bates *et al.*, 2015) and *LmerTest* (Kuznetsova *et al.*, 2017), and were optimized using the *bobyqa* optimizer.

*Linear relationship*

```
Model = lmer(day of first flower ~ latitude
        + (1 | year of observation),
        data = data,
        control=lmerControl(optimizer="bobyqa",optCtrl=list(maxfun = 100000)))
```

*Polynomial relationship*

```
Model = lmer(day of first flower ~ poly(latitude, degree=2)
        + (1 | year of observation),
        data = data,
        control=lmerControl(optimizer="bobyqa",optCtrl=list(maxfun = 100000)))
```

*Piecewise relationship*

This third model was parametrized using *nlme* (Pinheiro *et al.*, 2023) similarly to the first model and the estimation of the piecewise relationship is parametrized using *segmented* (Muggeo *et al.*, 2014). Here *nlme* was used because the output of *LmerTest* is not supported by *segmented.* The starting point value for the breakpoint estimation (psi) was the latitude of the minimum predicted day of first flower in the polynomial model.

Model = lme(day of first flower ~ latitude, random=~1| year of observation, data=data)

Segmented.Model = segmented.lme(Model, ~latitude, random=list(Year of observation = pdDiag(~1+latitude+U+G0)), psi=36.643)

**S2b: Parametrization of the hierarchical mixed-effect models testing for the effect of latitude and the predicted breakpoint on the day of first flower and climate variables.**

The best model out of the ones tested in Methods S2a identified a breakpoint at 34.98°N. The categorical fixed effect *breakpoint* was included in the model to depict the latitudinal position of observations relative to the identified breakpoint (< 34.98°N = 0, > 34.98°N = 1). Models were parametrized using the R packages *lme4* and *LmerTest* and were optimized using the *bobyqa* optimizer.

```
Model = lmer(dependent variable ~ latitude * breakpoint
        + (1 | Year of observation),
        data = data,
        control=lmerControl(optimizer="bobyqa",optCtrl=list(maxfun = 100000)))
```

**Methods S3: Raising of *Campanula americana* plants and seed rearing in Experiment 1.**

*Plant growth:*

Cohort 1 included 20 populations and Cohort 2 25 populations distributed across latitude. We sowed five seeds for each seed family in each population in the first cohort, two seeds per family per population for the second (one seed if few available, Table S3), in each of two pots filled with a 3:1 mixture of potting mix (LM-111 All Purpose Mix, Lambert, Rivière-Ouelle QC, Canada) and turface. Pots were randomly distributed across 128-cell propagation trays. Germination occurred in growth chambers under 12h light, 21°C day, 14°C night, 60% RH, for four weeks in Cohort 1, five in Cohort 2. Pots were watered daily for a week, then every other day. Trays were randomized within growth chambers twice a week. At the end of the germination, seedlings were randomly thinned to one per pot and plants were fertilized once (13 mg/L, Jack's Professional 15-5-15 + CaMg LX Fertilizer, JR Peters Inc., Allentown PA). Seedlings were then vernalized to simulate winter (4°C, low light for 12h per day), for six weeks in the first cohort and seven weeks in the second cohort. Half of the replicates were retained in each cohort after vernalization.

After vernalization, plants were moved to a greenhouse with daylength extended to 16h light with artificial light (600 µmol/$m^2$/s) to induce flowering. Initial temperature of 23°C day / 19°C night (+/- 5°C) was increased after two months to daytime temperature of 25°C and plants were shaded to simulate a light environment of an open forest. These temperatures reflected average summer temperatures across the range (Table S2b). Plants were watered as needed. Fertilizer was applied at 1000ppm (N) every two days for two months, then one to two times a week. Plants were treated with fungicide (two times before and two times after vernalization, Banrot 40 WP, Everris NA Inc., Dublin, OH; 0.9 g/L) to avoid root rot. During flowering, plants were sprayed with pesticide once a week to reduce damage by thrips (neem oil and liquid soap 1:1 and diluted in water at 13ml/L, and spinosad diluted in water at 16.5ml/L).

*Generation of laboratory-reared seed families:*

Plants raised in the first cohort were used to produce a greenhouse-reared generation of seeds for each population to be used for the second cohort. Each plant of the first cohort served as a pollen donor (father) and a pollen recipient (mother) in crosses to plants from the same population. Crosses were not reciprocal. Pollen-recipient flowers were emasculated in male phase to prevent selfing. Each cross was repeated three times, mature fruits were collected and stored in individual envelopes at 4°C under dry and dark conditions.

**Methods S4: Parametrization of the hierarchical mixed-effect models used in Experiments 1, 2, 3**

All models tested the effect of latitude and a categorical fixed effect (cohort, vernalization treatment, or common garden site), as well as their interactions, assuming either a linear or a quadratic relationship with latitude. For models with binary variables, we parametrized the models to use a logistic regression (*family = "binomial"*). For Experiment 2, the model also included the cross-random effect of the block (+ (1 | block), nested within a site) except for *transition to bolting* because it was analyzed based on population means within sites. Models were parametrized using the R packages *lme4* (Bates *et al.*, 2015) and *LmerTest* (Kuznetsova *et al.*, 2017), and were optimized using the *bobyqa* optimizer.

*Linear relationship with latitude*

```
Model = lmer(dependent variable ~ latitude * categorical effect
        + (1 | population / seed family within population),
        data = data,
        control=lmerControl(optimizer="bobyqa",optCtrl=list(maxfun = 100000)))
```

*Quadratic relationship with latitude*

```
Model = lmer(dependent variable ~ poly(latitude, degree=2) * categorical effect
        + (1 | population / seed family within population),
        data = data,
        control=lmerControl(optimizer="bobyqa",optCtrl=list(maxfun = 100000)))
```

**Supporting method S5: Estimation of first flowering time of *Campanula americana* in common gardens.**

*Estimation of time of first flowering protocol*

The time of first flower of plants grown in the common garden experiment was inferred from visits to each site two to three times early in flowering (Table S6b). At each visit, each plant was assigned one of 12 reproductive stages (-3 to 8), indicating its reproductive phenology at the time of the visit based on the condition of its buds, flowers and fruits (see Methods S5 Table below). These stages represent weekly intervals of reproductive phenology where stage 0 depicts plants that initiated flowering within a few days of the visit, negative stages depict plants that will flower *n* weeks after the visit, and positive stages represent plants that flowered *n* weeks before the visit. Estimates of weeks to flowering from these stages were based on observations in the greenhouse and prior field studies (personal observation). Occasionally, plants had characteristics of two successive stages and so were assigned the average of the two. Plants that had completed flowering and were dead (n=11) were assigned an additional stage (Stage 9, Method S5 Table).

We used the reproductive stage recorded at the visit closest to first flowering to infer the first flowering time for each plant. Little estimation is required for first flowering time for plants that flowered less than two weeks prior to data collection, or that will flower in the following week (i.e. between stages -1.5 and 2, Method S5 Table). These reproductive stages are easily identified and have a highly predictable relationship with flowering time. A total of 57% of the experimental plants were observed in these stages and hence the time of first flower was determined with little estimation. The reproductive stage recorded at the second and third visits was compared to each plant's predicted reproductive stage, given the stage recorded at the previous visit and the time elapsed between visits. The predicted reproductive stages were generally concordant with the recorded stage, with only 28 plants out of 2136 showing discrepancies. This indicates that our ability to predict plant progression through reproductive stages is robust and lends support to our method of predicting time of first flowering from reproductive stage.

*Evaluation of the accuracy of flowering time estimation*

Estimating the first flowering time from reproductive phenology has an increased chance of error for plants that flowered three or more weeks before a site was visited, or two or more weeks after a visit. We conducted several tests to assess the extent to which this estimation error may influence results. In a first "Reduced Estimation dataset," we tested the consequences of assigning successively earlier first flowering times for plants at later reproductive stages at the first visit by assigning all of these "early" flowering plants the same (early) first flowering time. Specifically, we took the conservative approach of assigning all plants with a reproductive stage greater than 2 to a reproductive stage of 3 and therefore the same flowering time. In a second "No Estimation dataset," we only included plants with a reproductive phenology between -1.5 and 2, and therefore there is little estimation of first flowering time (57% of original data). We analyzed these two datasets using the same methods as the full dataset but dropped populations where fewer than 5 individuals remained per site in the No Estimation dataset.

Analysis of the Reduced Estimation and No Estimation datasets yielded very similar results to the full dataset for three of the gardens (see below, Method S5 Figure). However, first flowering time in the southernmost garden was later in the Reduced Estimation and No Estimation datasets relative to the full data set. This is due to reducing information, either by removing plants that flowered substantially before our first visit, or assigning them a single first flowering time, rather than an estimated range of times. Most of these early flowering plants belonged to the earliest flowering populations in the other sites. In total, this suggests our estimates of the time of first flowering are likely to be fairly close to the actual first flowering time. The success of our estimation procedure is likely in part because previous observations of reproductive phenology in these sites were used to time our visits. Overall, our estimation procedure appears robust, and we therefore report the results of the full dataset (Fig. 3).

**Methods S5 Table: Estimation of time of first flowering in the common garden experiment based on reproductive stage.**

| Reproductive stage | Criteria |
|---|---|
| -3 | Apical meristem transitioned from producing only leaves to producing buds |
| -2 | Buds clearly visible, small and green |
| -1 | Large and purple buds, no open flowers (Fig. S2a) |
| 0 | First flower open, no dried flower or fruits (Fig. S2b) |
| 1 | First flowers dry but petals still colored, no fruits (Fig. S2c) |
| 2 | Many dried flowers, petals of some dry flowers brown, fruits not enlarged, many small green buds |
| 3 | Some fruits enlarged but not mature (green), many open and dry flowers, many small green buds |
| 4 | Some fruits mature (yellow to light brown), many open and dry flowers, few small green buds |
| 5 | Many fruits mature and some dehisced, buds still present |
| 6 | Many fruits dehisced, flowers present but no buds |
| 7 | Many fruits dehisced, few flowers |
| 8 | Many fruits dehisced, no flowers |
| 9 | Plant has reproduced and is completely dry |

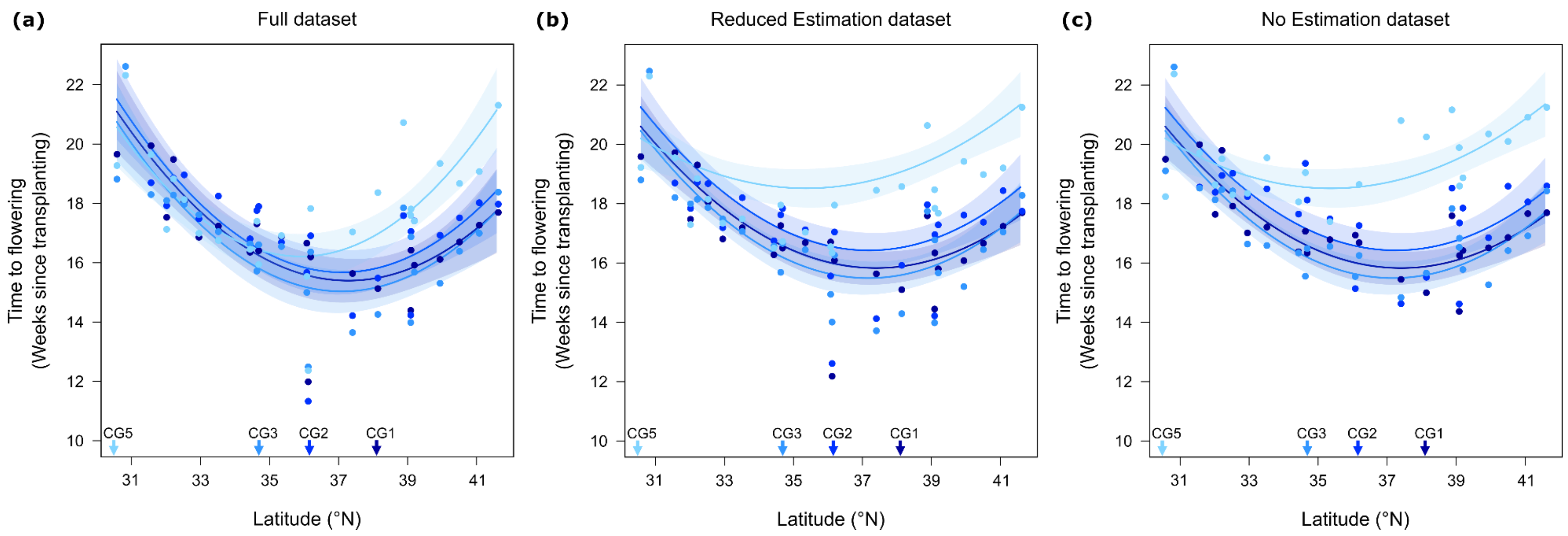


**Methods S5 Figure: Comparison of pattern in time to first flowering between the full dataset (a), Reduced Estimation dataset (b) and No Estimation dataset (c).** In the Reduced Estimation dataset, plants at later reproductive stages were assigned a uniform early first flowering time instead of estimating time of first flowering, while the No Estimation dataset included only plants with little estimation of phenology. Mean time to first flowering since transplant for populations (dots) sampled across the range and raised in four common gardens across a latitudinal gradient. Colors distinguish between the different gardens, and their latitude is indicated with arrows along the x-axis. Lines represent the significant model-predicted relationship for each garden between traits estimated at the individual level and home latitude of the population, with the 95% confidence interval indicated by shading. Note that panel (a) replicates Fig. 3a in the main text and is included here for ease of comparison.
